# The intrarenal renin-angiotensin system in hypertension: Insights from mathematical modelling

**DOI:** 10.1101/2021.12.14.472639

**Authors:** Delaney Smith, Anita Layton

**Affiliations:** Department of Applied Mathematics, University of Waterloo, 200 University Ave, Waterloo, N2L 3G1, ON, Canada; Cheriton School of Computer Science, University of Waterloo, 200 University Ave, Waterloo, N2L 3G1, ON, Canada; Department of Biology, University of Waterloo, 200 University Ave, Waterloo, N2L 3G1, ON, Canada; School of Pharmacy, University of Waterloo, 200 University Ave, Waterloo, N2L 3G1, ON, Canada

**Author notes:** Contributing authors.

**Keywords:** renin-angiotensin system, angiotensin II, hypertension, angiotensin II infusion, kidney, blood pressure regulation

## Abstract

The renin-angiotensin system (RAS) plays a pivotal role in the maintenance of volume homeostasis and blood pressure. In addition to the well-studied systemic RAS, local RAS have been documented in various tissues, including the kidney. Given the role of the intrarenal RAS in the pathogenesis of hypertension, a role established via various pharmacologic and genetic studies, substantial efforts have been made to unravel the processes that govern intrarenal RAS activity. In particular, several mechanisms have been proposed to explain the rise in intrarenal angiotensin II (Ang II) that accompanies Ang II infusion, including increased angiotensin type 1 receptor (AT1R)-mediated uptake of Ang II and enhanced intrarenal Ang II production. However, experimentally isolating their contribution to the intrarenal accumulation of Ang II in Ang II–induced hypertension is challenging, given that they are fundamentally connected. Computational modelling is advantageous because the feedback underlying each mechanism can removed and the effect on intrarenal Ang II can be studied. In this work, the mechanisms governing the intrarenal accumulation of Ang II during Ang II infusion experiments are delineated and the role of the intrarenal RAS in Ang II-induced hypertension is studied. To accomplish this, a compartmental ODE model of the systemic and intrarenal RAS is developed and Ang II infusion experiments are simulated. Simulations indicate that AT1Rmediated uptake of Ang II is the primary mechanism by which Ang II accumulates in the kidney during Ang II infusion. Enhanced local Ang II production is unnecessary. The results demonstrate the role of the intrarenal RAS in the pathogenesis of Ang II-induced hypertension and consequently, clinical hypertension associated with an overactive RAS.

## 1 Introduction

Hypertension is a common, highly complex condition that promotes risk for other diseases of the vasculature. Although the underlying causes of most cases of hypertension are unknown and likely multifactorial, antihypertensive therapies targeting the renin-angiotensin system (RAS) are highly effective in reducing elevated blood pressure (Ibrahim, 2006), due to the long-established role of the RAS in blood pressure regulation (Fyhrquist et al, 1995; Sparks et al, 2014). Angiotensin II (Ang II), the primary bio-active product of the RAS, increases blood pressure by inducing vasoconstriction, stimulating sympathetic nervous activity, increasing aldosterone production, and stimulating sodium reabsorption in the kidney (Fyhrquist et al, 1995; Sparks et al, 2014). Given the intrarenal actions of Ang II and the discovery that the kidney not only expresses, but independently regulates all components of the RAS (Kobori et al, 2007; Yim and Yoo, 2008; Li et al, 2019), the significance of the local intrarenal RAS to the pathology and development of hypertension has recently come into focus.

Ang II is produced systemically from the substrate angiotensinogen (AGT) following its release from the liver and its subsequent cleavage by various enzymes (Yim and Yoo, 2008; Nishiyama and Kobori, 2018). Renin is the first enzyme involved in the cascade, turning AGT into Ang I upon its secretion from the juxtamedullary apparatus of the kidney (Yim and Yoo, 2008; Nishiyama and Kobori, 2018). Chymase and angiotensin-converting enzyme (ACE) subsequently convert Ang I into Ang II, which exerts its effects upon binding to angiotensin type 1 and type 2 receptors (AT1Rs and AT2Rs, respectively) (Yim and Yoo, 2008). All hypertensive actions of Ang II, both systemic and intrarenal, are mediated by AT1R–Ang II binding (Navar et al, 2011; Crowley et al, 2006).

To prevent dis-regulation of the systemic RAS, renin secretion is moderated by Ang II itself via an AT1R–dependent mechanism. Indeed, renin secretion is inhibited (activated) by increased (decreased) Ang II levels to prevent the system’s over (under)-activation (Leete et al, 2018). However, given that the kidney independently regulates the production of many constituents of the RAS (Prieto-Carrasquero et al, 2004; Navar et al, 2011; Gonzalez and Prieto, 2015; Gonzalez et al, 2011; Navar et al, 2003; Kobori et al, 2001; Gonzalez-Villalobos et al, 2008; Zhuo et al, 2002, 1999; Cheng et al, 1995; Nishiyama and Kobori, 2018), systemic and intrarenal peptide concentrations can become decoupled. This disconnect is observed in cases of hypertension induced by Ang II infusion, a common protocol used to induce hypertension in preclinical models. In Ang II–induced hypertension, there is a progressive rise in intrarenal Ang II that cannot be explained on the basis of equilibration with plasma Ang II concentrations ([Ang II]) (Zou et al, 1996; Ellis et al, 2012; Gonzalez-Villalobos et al, 2008; Li et al, 2019).

Several mechanisms have been proposed to explain the renal accumulation of Ang II in Ang II-induced hypertensive rats, including: (i) enhanced AT1R– mediated uptake of circulating Ang II and (ii) increased intrarenal endogenous Ang II production (Kobori et al, 2007; Navar et al, 2011; Shao et al, 2009, 2010; Van Kats et al, 2001; Zhuo et al, 2002; Kobori et al, 2001, 2004; Ferrão et al, 2014; Gonzalez-Villalobos et al, 2008; Li et al, 2018b; Shao et al, 2009; Zou et al, 1998). In mechanism (i), circulating Ang II enters the kidney and binds to AT1Rs on tubular epithelial cells. The Ang II–AT1R complexes are then actively internalized (Hunyady et al, 2000) into intracellular compartments where the Ang II is protected from degradation (Zhuo et al, 2002; Navar et al, 2003; van Kats et al, 1997). It is hypothesized that the Ang II-dependent upregulation of AT1R expression in proximal tubule cells (Zhuo et al, 2002, 1999; Cheng et al, 1995) facilitates this effect. In mechanism (ii), the endogenous production of Ang II in the luminal fluid is thought to be increased as a result of Ang II-dependent positive feedback on proximal tubule AGT (Navar et al, 2003, 2011; Kobori et al, 2001; Gonzalez-Villalobos et al, 2008; Kobori et al, 2004; Schunkert et al, 1992) and collecting duct renin production (Prieto-Carrasquero et al, 2004; Navar et al, 2011; Gonzalez and Prieto, 2015; Gonzalez et al, 2011).

While these hypotheses are well-founded, isolating the contribution of each mechanism to the accumulation of Ang II in the kidney in Ang II–induced hypertension poses a significant experimental challenge because they are fundamentally linked. Computational modelling is advantageous in this regard, because once a model is developed that incorporates these systems, each feedback function can be individually turned on/off and the impact on the total and local intrarenal [Ang II] can be studied. Two computational models of the intrarenal RAS have been developed previously (Schalekamp and Danser, 2006; Lo et al, 2011). However, neither considers intrarenal positive feedback. The first model (Schalekamp and Danser, 2006) was designed to estimate the intrarenal distribution of Ang II and AT1/2Rs. However, it does not consider any temporal dynamics and thus can only be used to study the system’s behaviour at steady state. While the second model (Lo et al, 2011) does consider the rate of change of key RAS peptides, it does not differentiate between the intracellular and extracellular compartments of the kidney. Therefore, it cannot be used study the process of AT1R-mediated uptake of circulating Ang II (mechanism (i)) in detail. For these reasons, the existing models would be ineffectual in studying the contribution of the intrarenal RAS to the pathogenesis of Ang II-induced hypertension.

In this work, we aim to delineate the mechanisms that mediate intrarenal Ang II accumulation during Ang II infusion and consequently, gain insight into the role of the intrarenal RAS in the development of Ang II-induced hypertension. To do so, we develop a novel computational model of the intrarenal RAS that considers temporal dynamics, distinguishes between the intracellular and extracellular regions of various intrarenal compartments, and incorporates intrarenal (and systemic) positive feedback. Formalism’s to simulate both subcutaneous and intravenous Ang II infusion experiments are also derived to facilitate the study of Ang II–induced hypertension. After fitting the model parameters to both steady state and Ang II infusion data, the model is validated and used to make predictions on the key mechanisms contributing to the rise in intrarenal [Ang II] during Ang II infusion. The robustness of the predictions is also quantified via a local parametric sensitivity analysis. The results presented provide novel insight into the role of the intrarenal RAS in the development of Ang II-induced hypertension and accordingly, any form of hypertension associated with an overactive RAS.

## 2 Methods

### 2.1 Intrarenal RAS model

The intrarenal model considers Ang I and II dynamics across four tissue compartments: the glomerular compartment, the peritubular compartment, the tubular compartment, and the vasculature compartment. The vasculature compartment comprises both the lymphatic vasculature and the blood vasculature. The other compartments are subdivided into extracellular and intracellular (endosomal) regions, connected to one another via AT1R–bound Ang II internalization. Since Ang I does not bind to AT1Rs, it is assumed to be restricted to the extracellular regions. This assumption is consistent with data from (Imig et al, 1999), which indicates that intracellular Ang I comprises a negligible portion of total renal Ang I (4%).

The luminal fluid constitutes the extracellular region of the tubular compartment and the interstitium comprises the extracellular region of both the glomerular and peritubular compartments. The glomerular and peritubular interstitium are separated at the level of the glomerular vascular pole, such that the glomerular interstitium represents the mesangium. The intracellular region of the glomerular compartment comprises of mesangial cells, whereas that of the tubular and peritubular compartments comprises of epithelial cells. The intracellular tubular and peritubular regions differ in that the former represents apically-derived endosomes, while the latter represents basolaterally-derived endosomes. A visual representation of each compartment as they appear in the kidney is presented in Figure Ia, with the physiological processes connecting the compartments outlined in Figure Ib.

**Fig. I:**
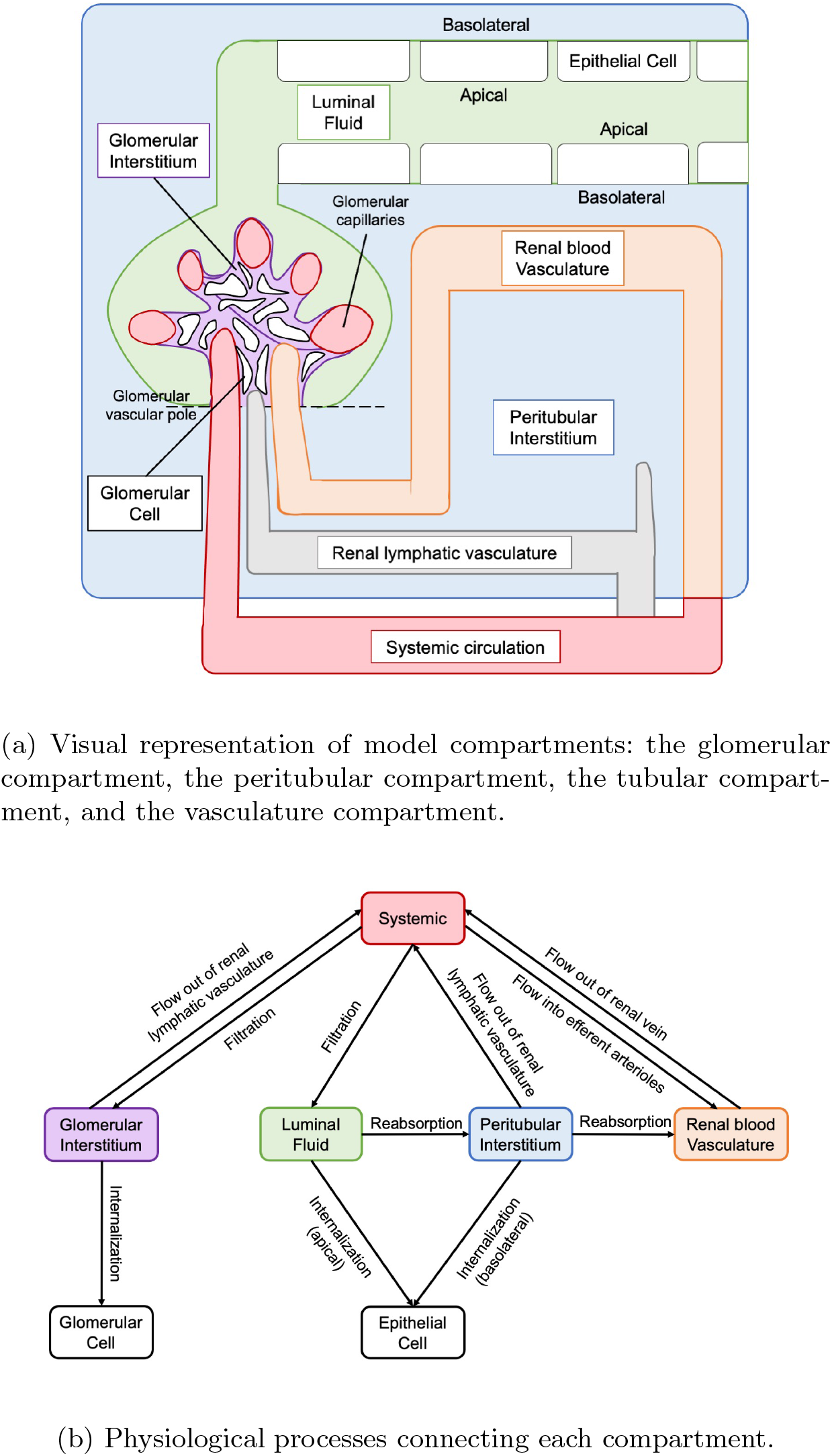
Model schematics

The mathematical formalisms used to model the aforementioned processes are described in detail below. The structure of the equations describing Ang I, Ang II, AT1R–bound Ang II, and AT1R dynamics are similar, albeit not exactly the same, across all renal compartments. Therefore, we introduce general equations describing each variable in the intracellular (*L* = *Cell*), membrane–bound (*L* = *Memb*), and extracellular (*L* = *Ext*) regions of each compartment *C* (*C* = *Gl, Tb*,, *Pt, Pv*) in section 2.1.1, before detailing the specifics of each compartment in sections 2.1.2 – 2.1.5.

#### 2.1.1 General model equations

The Ang I and Ang II dynamics in renal sub-compartment *C*_*L*_ (*L* = *Ext, Cell*) can be described by the following equations:

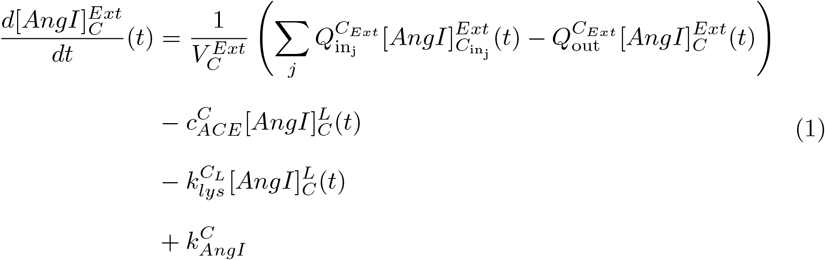

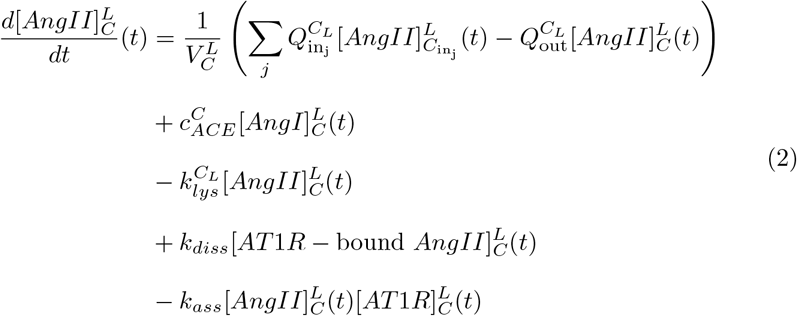

The first line of each equation represents the balance between the *j* fluxes into sub-compartment *C*_*L*_ and the flux out of sub-compartment *C*_*L*_. The second line represents the conversion of Ang I to Ang II by local ACE activity with rate 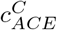. The third line represents peptide degradation with rate 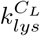. Intracellular Ang II is assumed to be degraded by lysosomes and peptide X (*X* = *AngI, AngII*) in the renal blood vasculature is assumed to decay according to its half life *h*_*X*_ :

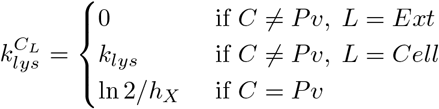

The rest of the expressions used to describe the dynamics of Ang I and Ang II differ. Indeed, as described by the last line of Eq. 1, Ang I is endogenously produced from local renin activity at an assumed constant rate of 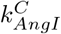. In contrast, the fourth and fifth lines of Eq. 2 represent the unbinding and binding of Ang II to AT1Rs with rates *k*_*diss*_, and *k*_*ass*_, respectively. Finally, since Ang II is assumed to enter the intracellular sub-compartments *C*_*Cell*_ via AT1R–bound Ang II internalization only, we set 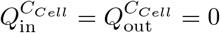 in all compartments *C*.

The equation describing membrane–bound (*L* = *Memb*) and intracellular (*L* = *Cell*) AT1R–bound Ang II dynamics in each compartment *C* is given by:

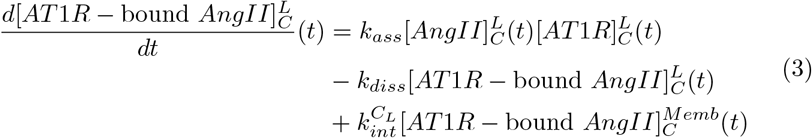

The first line represents the binding of Ang II to AT1Rs with rate *k*_*ass*_, the second line represents the unbinding of Ang II from AT1Rs with rate *k*_*diss*_, and the third line represents membrane-bound AT1R–bound Ang II internalization. A simplified model of AT1R binding that does not consider membrane-bound AT1R-Ang II complex internalization was implemented in the renal blood vasculature (and systemic circulation, see section 2.2), due to the substantial uncertainties in key variables such as the volume of such an intracellular compartment and the concentration of intracellular endothelial Ang II. Hence:

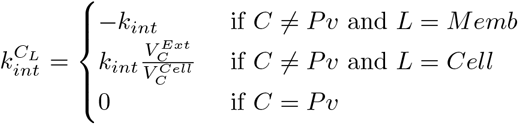

The equation describing intracellular AT1R dynamics in the glomerular, tubular, and peritubular compartments (*C* = *Gl, T b, Pt*) is given by:

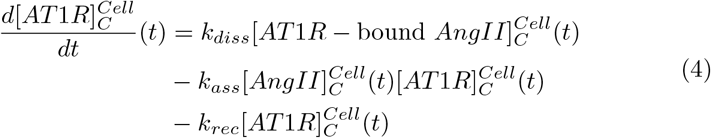

The first line represents the unbinding of Ang II from AT1Rs with rate *k*_*diss*_, the second line represents the binding of Ang II to AT1Rs with rate *k*_*ass*_, and the third line represents the recycling of internalized AT1Rs back to the cell membrane. It is assumed that the AT1Rs are recycled back to the same membrane (basolateral or apical) and thus compartment (peritubular or tubular) from which they originate in tubular epithelial cells.

Finally, the equations describing the concentration of membrane-bound AT1Rs, 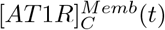, differ across renal compartments. A compartment-specific derivation of these equations as well as a description of those outlined above is given in sections 2.1.2–2.1.5 below.

#### 2.1.2 Glomerular compartment

A small fraction of the renal plasma flow enters the glomerular interstitium (denoted *ϕ*_*L*_) (Lemley et al, 1992), contributing to an influx of circulating Ang I and II to this sub-compartment. This influx is balanced by lymphatic drainage with flow rate *ϕ*_*L*_. Given the lack of data surrounding ACE activity and the concentration of Ang I and II in the glomerulus, we set 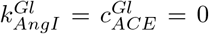. Furthermore, to derive the equation for membrane-bound AT1R dynamics, we assume that the total amount of AT1Rs in the glomerular compartment is conserved, such that:

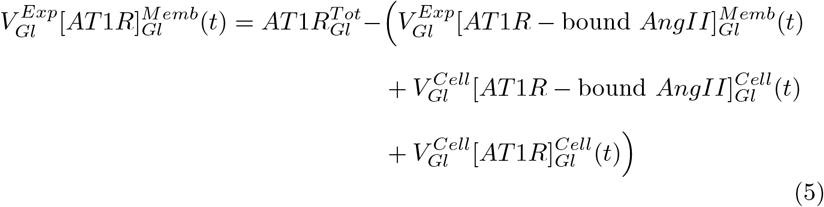

#### 2.1.3 Tubular compartment

Ang I and II are filtered from the systemic circulation with rate *ϕ*_*GFR*_, reabsorbed across the tubular epithelium at a rate proportional to fluid flow *ϕ*_*Pt*_, and cleared in the urine with rate *ϕ*_*U*_. We assume that only a fraction *S*_*Pt*_ ∈ [0, 1] of angiotensin is transported across the the tubular epithelium alongside fluid flow. Conservation of flow is assumed such that *ϕ*_*Pt*_ = *ϕ*_*GFR*_ − *ϕ*_*U*_. Furthermore, as in the glomerular compartment, we assume that the total amount of tubular AT1Rs is conserved (*f*_*Tb*_ = 0, see below) in healthy steady state conditions. However, in conditions where local Ang II concentrations are elevated, AT1R expression has been observed to increase (*f*_*Tb*_ > 0) via an AT1R-dependent mechanism (Zhuo et al, 2002, 1999; Cheng et al, 1995). This positive feedback loop (assumed linear for simplicity) permits the accumulation of Ang II in intracellular endosomes in cases of Ang II–induced hypertension (Zhuo et al, 2002). Given that similar feedback is included in the peritubular and circulating compartments (see below), we introduce a general description of the function *fb*_*C*_ here (*C* = *T b, P t, circ*(*AGT*), *circ*(*ACE*)), along with the equation describing tubular membrane-bound AT1R dynamics:

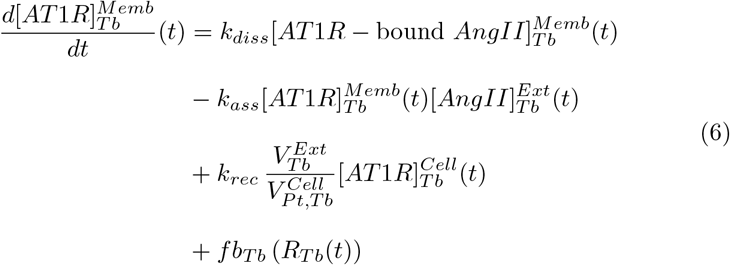

where

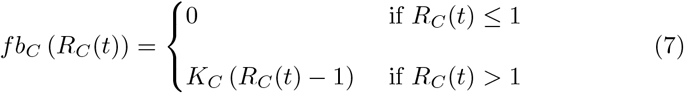

and

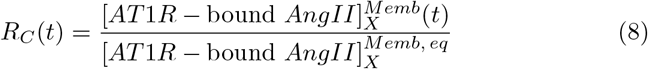

#### 2.1.4 Peritubular compartment

Ang I and Ang II are reabsorbed from the luminal fluid into the peritubular interstitium with rate *S*_*Pt*_*ϕ*_*Pt*_. Peritubular interstitial peptides are then drained via the lymphatic vasculature and reabsorbed into the renal blood vasculature at rates proportional to fluid flow *ϕ*_*L*_ and *ϕ*_*Pv*_, respectively. Conservation of flow is assumed, such that *ϕ*_*Pv*_ = *ϕ*_*GFR*_ − *ϕ*_*L*_ − *ϕ*_*U*_. Moreover, peritubular AT1Rs behave identically to tubular AT1Rs in that their total concentration is conserved in healthy steady state conditions (*fb*_*Pt*_ = 0), but up-regulated when local Ang II concentrations rise (*fb*_*Pt*_ > 0) (Zhuo et al, 2002, 1999; Cheng et al, 1995). This positive feedback (Eq. 7) is assumed to be driven by AT1R-signalling (Zhuo et al, 2002). Hence, the equation describing peritubular membrane-bound AT1R dynamics is given by:

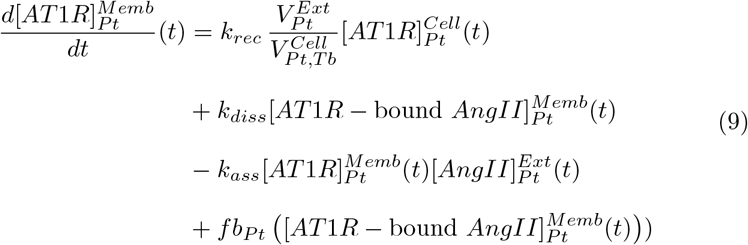

#### 2.1.5 Blood vasculature compartment

The amount of renal plasma flow that does not enter the nephron or glomerular interstitium, i.e. *ϕ*_*RPF*_ − *ϕ*_*L*_ − *ϕ*_*GFR*_, enters the efferent arterioles, bringing a proportional amount of circulating Ang I and II with it. Also impacting the rate of change of [Ang I] and [Ang II] at this level is re-absorption from the peritubular interstitium at a rate proportional to fluid flow *ϕ*_*Pv*_ and loss via the renal vein. Once again, flow is assumed to be conserved with concentrations of Ang I and II proportional to

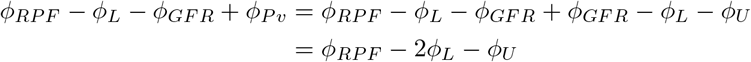

being returned to the systemic circulation.

As aforementioned, a simplified model of AT1R binding that does not consider membrane-bound AT1R-Ang II complex internalization was implemented in the blood vasculature. Instead, the total concentration of AT1Rs in this compartment 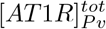 is assumed to remain constant:

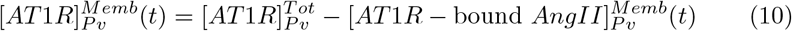

#### 2.1.6 Whole kidney concentrations

The amounts of Ang I and II in each renal compartment described above are summed to equal total renal concentrations in units of *fmol/g* kidney.

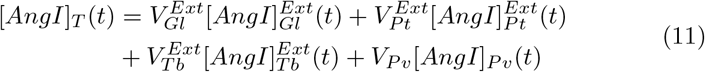

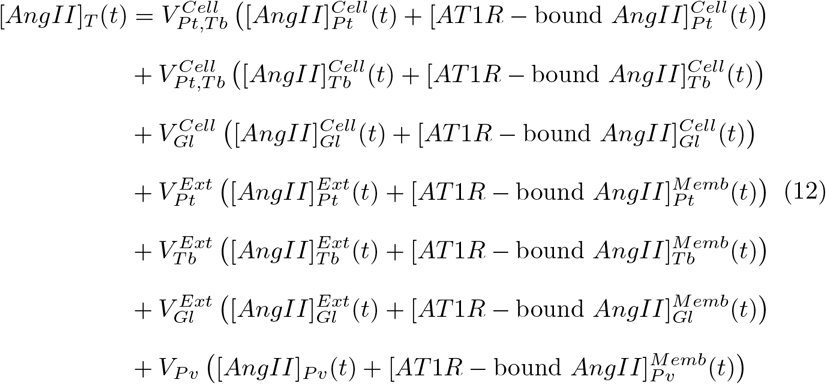

A summary of the parameters outlined in Eqs. 1 – 4 specific to each sub-compartment is given in Table II below. The complete set of intrarenal model equations is provided in Appendix A.1.

**Table I:**
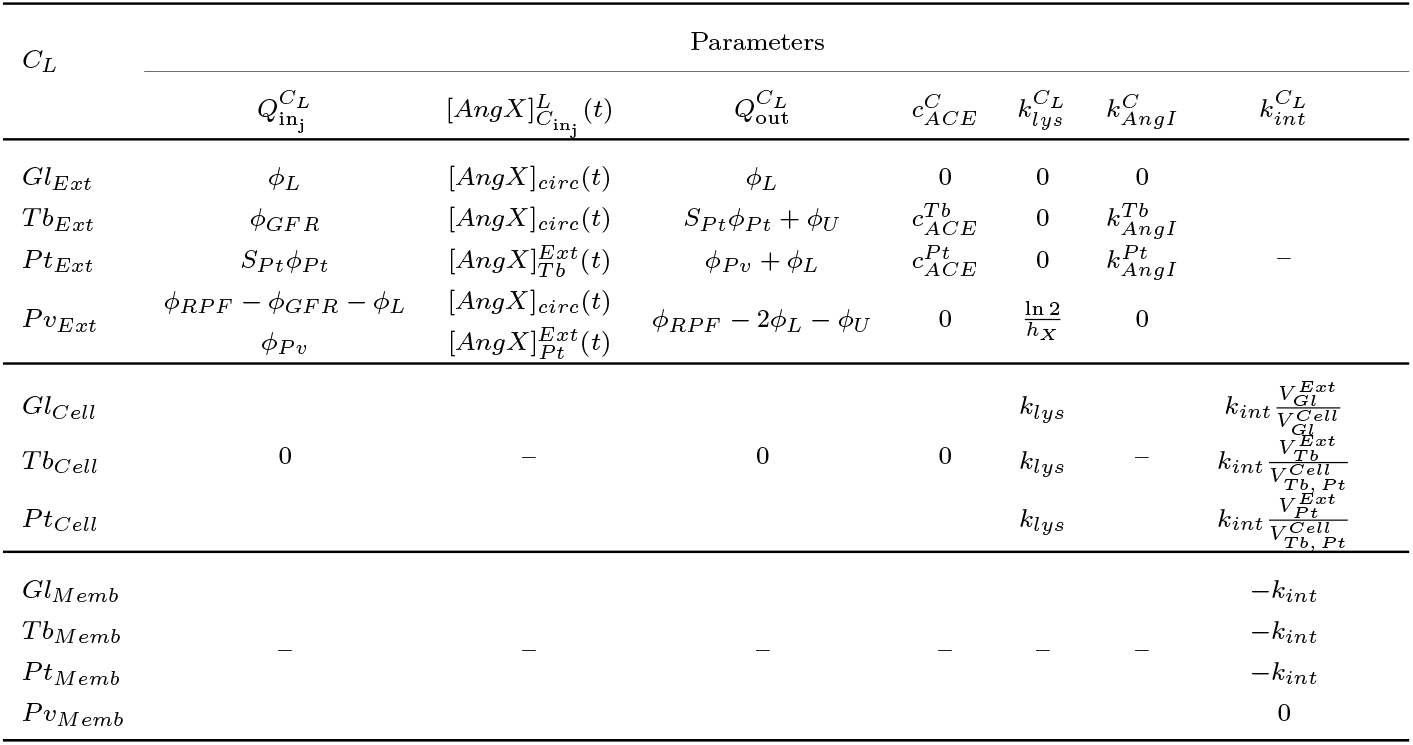
Sub-compartment-specific parameters corresponding to Eqs. 1–4.

**Table II:**
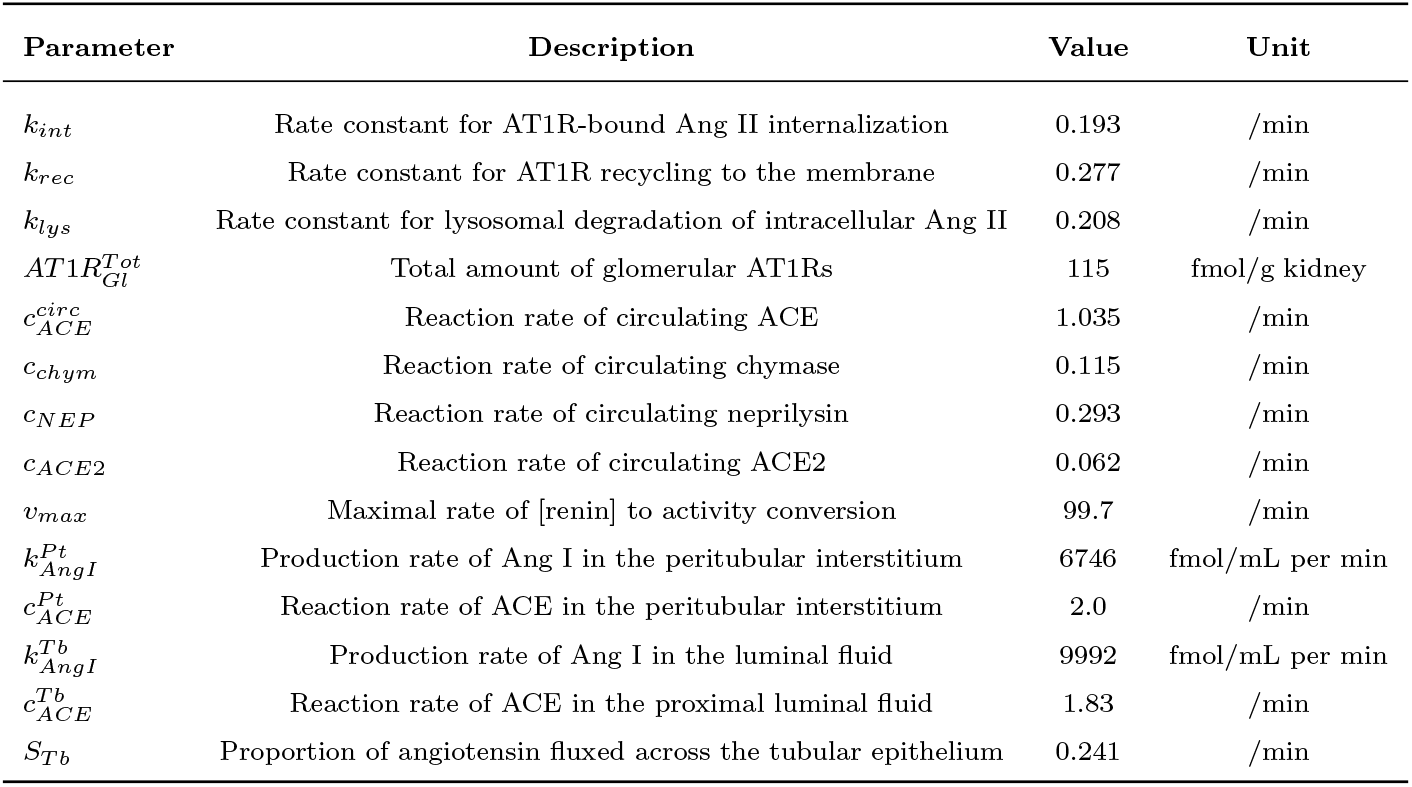
Baseline fitted parameters

| Parameter | Description | Value | Unit |
| --- | --- | --- | --- |
| $k_{int}$ | Rate constant for AT1R-bound Ang II internalization | 0.193 | /min |
| $k_{rec}$ | Rate constant for AT1R recycling to the membrane | 0.277 | /min |
| $k_{lys}$ | Rate constant for lysosomal degradation of intracellular Ang II | 0.208 | /min |
| $AT1R_{Gl}^{Tot}$ | Total amount of glomerular AT1Rs | 115 | fmol/g kidney |
| $c_{ACE}^{circ}$ | Reaction rate of circulating ACE | 1.035 | /min |
| $c_{chym}$ | Reaction rate of circulating chymase | 0.115 | /min |
| $c_{NEP}$ | Reaction rate of circulating neprilysin | 0.293 | /min |
| $c_{ACE2}$ | Reaction rate of circulating ACE2 | 0.062 | /min |
| $v_{max}$ | Maximal rate of [renin] to activity conversion | 99.7 | /min |
| $k_{AngI}^{Pt}$ | Production rate of Ang I in the peritubular interstitium | 6746 | fmol/mL per min |
| $c_{ACE}^{Pt}$ | Reaction rate of ACE in the peritubular interstitium | 2.0 | /min |
| $k_{AngI}^{Tb}$ | Production rate of Ang I in the luminal fluid | 9992 | fmol/mL per min |
| $c_{ACE}^{Tb}$ | Reaction rate of ACE in the proximal luminal fluid | 1.83 | /min |
| $S_{Tb}$ | Proportion of angiotensin fluxed across the tubular epithelium | 0.241 | /min |

### 2.2 Systemic RAS model

The systemic RAS model is based on those by Leete et al (2018) and Ahmed and Layton (2020). Modifications were made to couple the systemic model to the intrarenal model, as well as to the AT1R–Ang II binding and plasma renin activity formalisms.

The rate of change of the plasma AGT concentration ([*AGT* ]_*circ*_) is governed by its endogenous production at a constant rate *k*_*AGT*_, conversion to Ang I by renin, and degradation with a half-life *h*_*AGT*_. Moreover, elevated systemic [Ang II] has been observed to increase hepatic angiotensinogen production in a AT1R-dependent manner (Li and Brasier, 1996; Schunkert et al, 1992; Nakamura et al, 1990). This positive feedback is modelled by the addition of the function *fb*_*circ*(*AGT*)_ (Eq. 7) to the equation describing plasma [*AGT* ] dynamics:

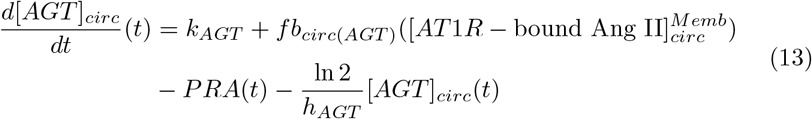

*PRA* denotes plasma renin activity. As in Leete et al (2018), *PRA* is assumed to follow Michaelis-menten kinetics, with the Michaelis constant *K*_*M*_ reflecting the concentration of *AGT* where *PRA* is half-maximal. However, similarly to the formulation by Ahmed and Layton (2020), the maximal activity is assumed to depend linearly on the plasma renin concentration *PRC*, with rate constant *v*_*max*_. In this way, sufficient concentrations of both renin and AGT are required to generate sufficient plasma renin activity:

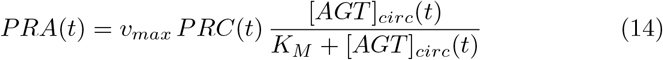

Renin is secreted from the juxtamedullary apparatus of the kidney at a basal rate *R*_*sec*_. This secretion rate is modified by the concentration of glomerular membrane-bound AT1R-Ang II complexes, 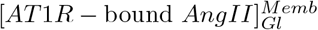, via the feedback function *ν*_*AT*1*R*_ (Eq. 16). Since the exponent *B*_*AT*1*R*_ is always positive, *ν*_*AT*1*R*_ drops below 1 to inhibit renin secretion when 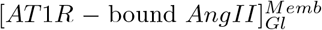 rises above its healthy steady state 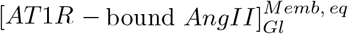, i.e. when their ratio *R*_*Gl*_ to rise above 1. The opposite effect is observed if 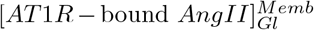 drops below its steady state, i.e. when *R*_*Gl*_ drops below 1. Also contributing to the rate of change of *PRC* is renin’s decay according to its half-life *h*_*renin*_, resulting in the equation:

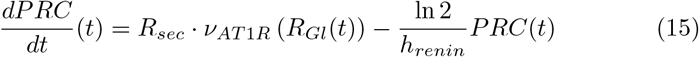

where

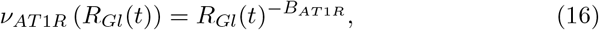

Plasma Ang I, *AngI*_*circ*_ decays with a half-life *h*_*AngI*_ and is converted into other forms by ACE, neprilysin (NEP), and chymase activity with rates 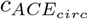, *c*_*NEP*_, and *c*_*chym*_, respectively. Systemic ACE activity is assumed to increase in an Ang II and AT1R-dependent manner via the positive feedback function *fb*_*circ*(*ACE*)_ (Eq. 7). This feedback was found to be required to replicate the decrease (increase) in plasma Ang I (endogenous Ang II) observed experimentally following Ang II infusion (see section 3.1) (Zou et al, 1996; Shao et al, 2009, 2010). Similar feedback has been documented in the kidney (Sadjadi et al, 2005; Koka et al, 2008). In addition, plasma Ang I enters the kidney via the renal artery. In doing so, an amount of Ang I proportional to *ϕ*_*RPF*_ gets distributed to the various renal compartments as described above. Following local renal modifications, Ang I from the renal blood vasculature and interstitium gets returned to the systemic circulation via the renal vein and lymphatic vasculature, respectively, with corresponding flow rates *ϕ*_*RPF*_ − 2*ϕ*_*L*_ − *ϕ*_*U*_ and *ϕ*_*L*_. In this way, fluid flow into and out of the kidney is conserved, differing only by a factor of *ϕ*_*U*_ which is assumed to be balanced by water intake:

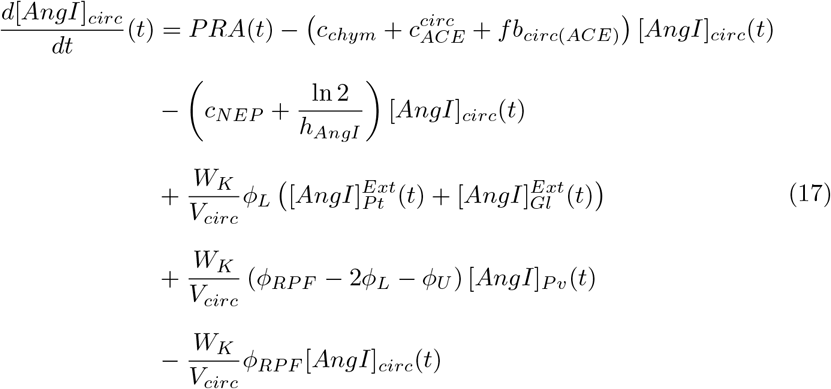

The rate of change of plasma [*Ang*(1 − 7)] ([*Ang*(1− 7)]_*circ*_) is governed by its production from Ang I by neprilysin and Ang II by ACE2 with respective rate constants *c*_*NEP*_ and *c*_*ACE*2_, as well as its decay according to its half-life *h*_*Ang*17_:

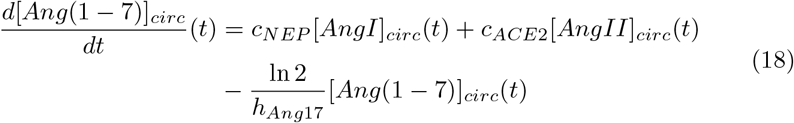

Systemic Ang II (*AngII*_*circ*_) is produced from Ang I by chymase and ACE activity, converted to *Ang*(1 − 7) by ACE2 activity, and decays with half-life *h*_*AngII*_. Identically to Ang I, systemic Ang II enters the kidney via the renal artery, undergoes local modifications, and is returned to the circulation via the renal vein and lymphatic vasculature. Unlike Ang I however, Ang II may bind to AT1Rs in the systemic vasculature endothelium, 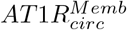 with rate *k*_*ass*_ to form membrane-bound AT1R-Ang II complexes, *AT*1*R* − bound 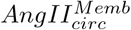. This binding is reversible, with dissociation rate *k*_*diss*_. As in the renal blood vasculature, the total concentration of systemic AT1Rs 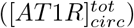 is assumed constant:

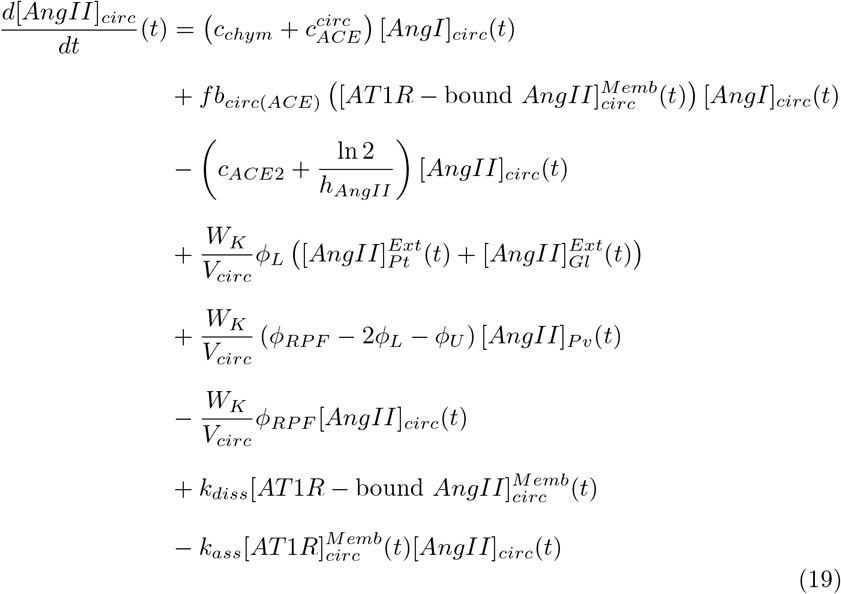

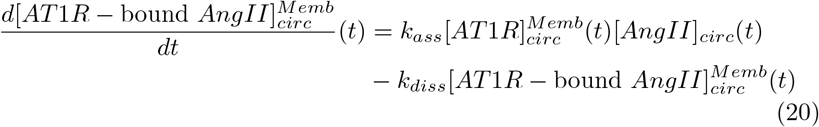

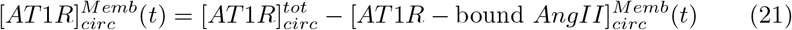

A schematic outlining the connections between all model variables is shown in Figure II.

**Fig. II:**
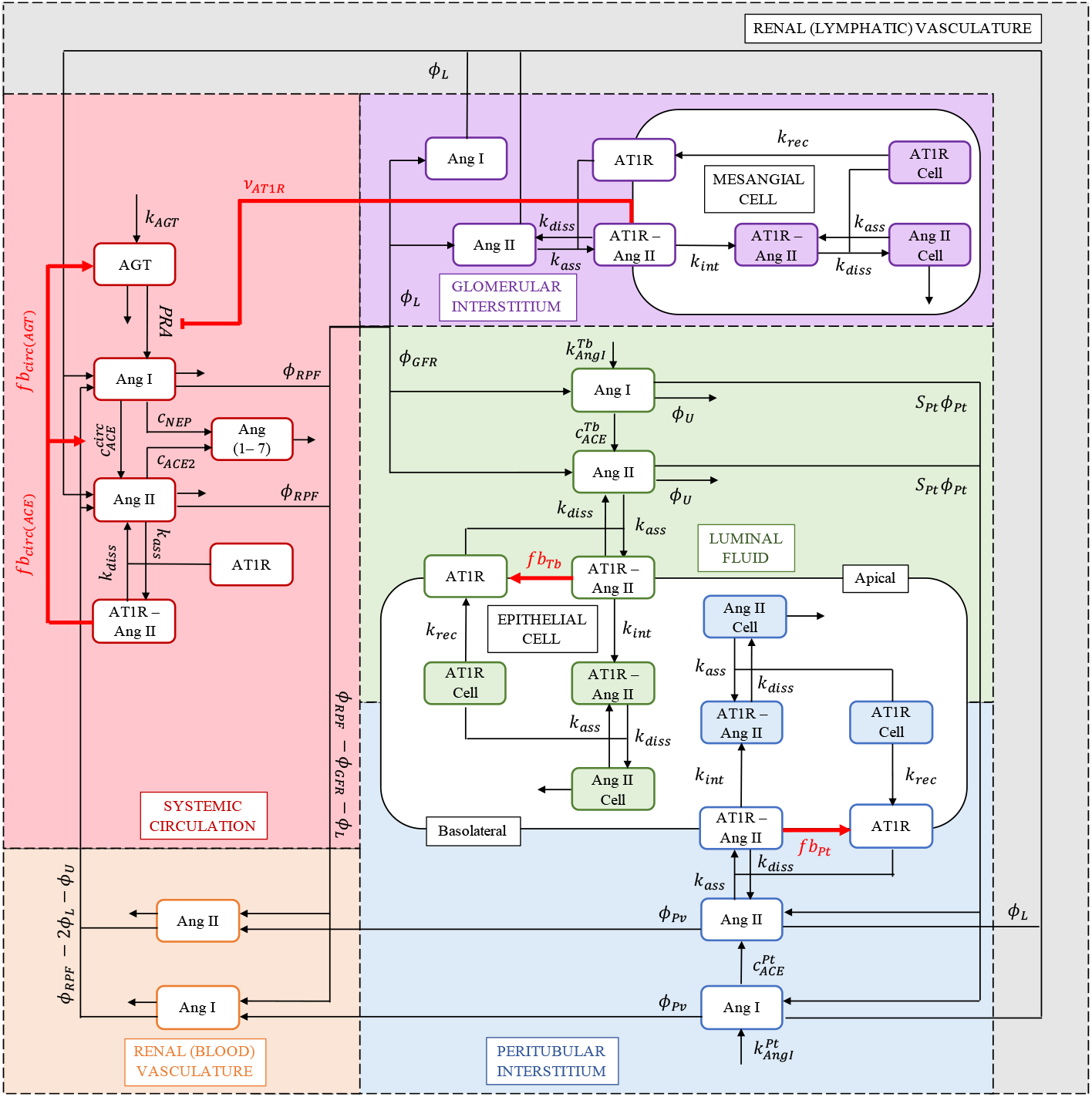
Schematic representation of the connections between model variables. Red connections indicate feedback.

To fit and validate the model and to investigate the changes to the intrarenal RAS that occur during the development of Ang II-induced hypertension, we simulate Ang II infusion experiments.

### 2.3 Simulating Ang II infusion experiments

Ang II can be infused intravenously (IV), or subcutaneously (SC) via osmotic mini-pump implantation. Each experimental technique is simulated using a different mathematical formalism.

During an IV infusion, Ang II enters the blood stream directly. Hence, it is simulated by adding the constant production term:

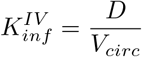

to the original equation describing plasma Ang II dynamics (Eq. 19). *D* is the dose of Ang II in units of fmol/min, and *V*_*circ*_ is the circulating plasma volume in mL.

During a SC infusion, Ang II enters the blood stream indirectly following re-absorption from the SC tissue. Hence, we must consider the amount of Ang II in this compartment, *AngII*_*SC*_. The rate of change of *AngII*_*SC*_ is governed by the exogenous infusion of Ang II into this compartment with dose *D* (fmol/min) and its re-absorption into the systemic circulation with rate *k*_*a*_ min^−1^:

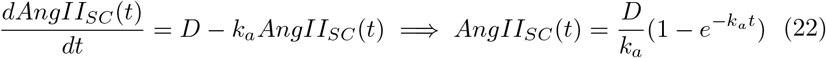

Therefore, SC Ang II infusion can be simulated by adding the following term to the original ODE describing [*AngII*]_*circ*_ (Eq. 19).

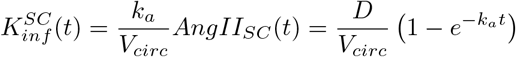

#### 2.3.1 Separating exogenous Ang II from endogenous Ang II

To better understand Ang II dynamics, we separately simulate in each compartment exogenously infused (*AngII* − *i*) and endogenously produced (*AngII* − *p*) Ang II, such that

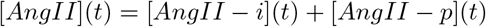

The equations describing exogenous and endogenous Ang II differ in their production terms: Infusion contributes to *AngII*− *i* levels, while enzyme activity contributes to *AngII* − *p* levels. In this way, the equations describing *AngII*− *p* are analogous to those described above in section 2.1.1 (Eqs. 2, 3) and the equations describing *AngII*− *i* only differ in the compartments where the endogenous production of Ang II occurs (systemic circulation, peritubular interstitium, and luminal fluid). Indeed, the infusion term 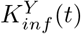 (*Y* = *SC, IV*) is added to the systemic circulation (Eq. 19) to simulate infusion and the terms describing chymase and/or ACE activity are removed in all 3 of the aforementioned compartments i.e. we set 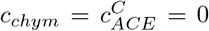 for *C* = *T b, P t, circ* (Eqs. 2, 19).

In the absence of an infusion, [*AngII*− *i*] = 0 in all compartments and [*AngII* − *p*] is the sole contributor to the total Ang II concentration, as expected.

### 2.4 Model parameter estimation

We divide the model parameters into two groups; i) those that impact the system’s steady state (baseline parameters), and ii) those that influence the system only when it is perturbed from this steady state (feedback parameters). These parameter groups are identified sequentially, as outlined below.

Many baseline parameters (Table AI) were derived from the literature, as described in Appendix A.2. To identify the remaining parameters in the baseline set *p*_0_ (Table II) and an appropriate steady state for the model variables *x*_0_ (Table III), we minimize the cost function

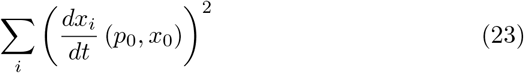

subject to the following conditions, where *x*_*i*_ represents variable *i* in the model:

1. Since AngII–AT1R–mediated endocytosis occurs with a half-life of 2 to 10 minutes (Thomas et al, 1996), the rate constant describing AngII−AT1R complex internalization, *k*_*int*_ (Eq. 3), was restricted to the interval [0.07, 0.35] /min (Schalekamp and Danser, 2006). The same upper bound (0.35/min) was assumed for the rate constants describing receptor recycling *k*_*rec*_ (Eqs. 4, 6, 9) and lysosomal degradation *k*_*lys*_ (Eq. 2).
2. The total amount of glomerular 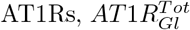 (Eq. 5), was fit to ensure that the total concentration of glomerular receptors was greater than the total concentration of tubular and peritubular receptors at baseline (Zhuo et al, 1998), but less than the upper bound of 1000*K*_*D*_ (Schalekamp and Danser, 2006). *K*_*D*_ = *k*_*diss*_*/k*_*ass*_ is the dissociation constant of Ang II from AT1Rs. The same value (1000*K*_*D*_) was used to bound the total concentration of AT1Rs in all other compartments.
3. The remaining parameters were fit to ensure that the steady state concentrations were within the range of the renal and systemic RAS concentrations that have been observed experimentally (Table IV).

**Table III:** Steady state concentrations of model variables

| Variable | Description | Steady State | Unit |
| --- | --- | --- | --- |
| Systemic compartment |  |  |  |
| $[AGT]_{circ}$ | Plasma [angiotensinogen] | 575965 | |
| $[AngI]_{circ}$ | Plasma [Ang I] | 65.3 | |
| $[AngII]_{circ}$ | Plasma [Ang II] | 43.4 | |
| $[AT1R - \text{bound } AngII]_{circ}^{Mem}$ | Membrane-bound [AT1R-Ang II complex] | 21679 | fmol/mL |
| $[AT1R]_{circ}^{Mem}$ | Membrane-bound free [AT1R] | 499821 | |
| $[Ang(1 - 7)]_{circ}$ | Plasma [Ang (1-7)] | 5.25 | |
| PRC | Plasma [renin] | 4.33 |  |
| PRA | Plasma renin activity | 73.6 | fmol/mL per min |
| Glomerular compartment |  |  |  |
| $[AngI]_{Gl}^{Ext}$ | Interstitial [Ang I] | 65.3 | |
| $[AngII]_{Gl}^{Ext}$ | Interstitial [Ang II] | 40.9 | |
| $[AT1R - \text{bound } AngII]_{Gl}^{Mem}$ | Glomerular membrane-bound [AT1R-Ang II complex] | 262.4 | fmol/mL |
| $[AT1R - \text{bound } AngII]_{Gl}^{Cell}$ | Intracellular [AT1R-Ang II complex] | 2154 | |
| $[AngII]_{Gl}^{Cell}$ | Free intracellular [Ang II] | 243.4 | |
| $[AT1R]_{Gl}^{Mem}$ | Free membrane-bound [AT1R] | 57927 | |
| $[AT1R]_{Gl}^{Cell}$ | Free intracellular [AT1R] | 182.6 | |
| Peritubular compartment |  |  |  |
| $[AngI]_{Pt}^{Ext}$ | Interstitial [Ang I] | 454 | |
| $[AngII]_{Pt}^{Ext}$ | Interstitial [Ang II] | 189 | |
| $[AT1R - \text{bound } AngII]_{Pt}^{Mem}$ | Basolateral membrane-bound [AT1R-Ang II complex] | 88.6 | fmol/mL |
| $[AT1R - \text{bound } AngII]_{Pt}^{Cell}$ | Basolateral intracellular [AT1R-AngII complex] | 57.2 | |
| $[AngII]_{Pt}^{Cell}$ | Free basolateral intracellular [Ang II] | 6.60 | |
| $[AT1R]_{Pt}^{Mem}$ | Free basolateral membrane-bound [AT1R] | 4232 | |
| $[AT1R]_{Pt}^{Cell}$ | Free basolateral intracellular [AT1R] | 4.95 | |
| Tubular compartment |  |  |  |
| $[AngI]_{Tb}^{Ext}$ | Luminal fluid [Ang I] | 1634 | |
| $[AngII]_{Tb}^{Ext}$ | luminal fluid [Ang II] | 757 | |
| $[AT1R - \text{bound } AngII]_{Tb}^{Mem}$ | Apical membrane-bound [AT1R-Ang II complex] | 26.9 | fmol/mL |
| $[AT1R - \text{bound } AngII]_{Tb}^{Cell}$ | Apical intracellular [AT1R-Ang II complex] | 75.1 | |
| $[AngII]_{Tb}^{Cell}$ | Free apical intracellular [Ang II] | 8.67 | |
| $[AT1R]_{Tb}^{Mem}$ | Free apical membrane-bound [AT1R] | 321 | |
| $[AT1R]_{Tb}^{Cell}$ | Free apical intracellular [AT1R] | 6.50 | |
| Renal blood vasculature compartment |  |  |  |
| $[AngI]_{Pv}$ | Renal plasma [Ang I] | 163.8 | fmol/mL |
| $[AngII]_{Pv}$ | Renal plasma [Ang II] | 79.1 | |
| $[AT1R - \text{bound } AngII]_{Pv}^{Mem}$ | Membrane-bound [AT1R-AngII complex] | 36.7 | |
| $[AT1R]_{Pv}^{Mem}$ | Free membrane-bound [AT1R] | 463 | |
| Whole kidney |  |  |  |
| $[AngI]_T$ | Whole kidney [Ang I] | 191 | fmol/g kidney |
| $[AngII]_T$ | Whole kidney [Ang II] | 145 | |

**Table IV:**
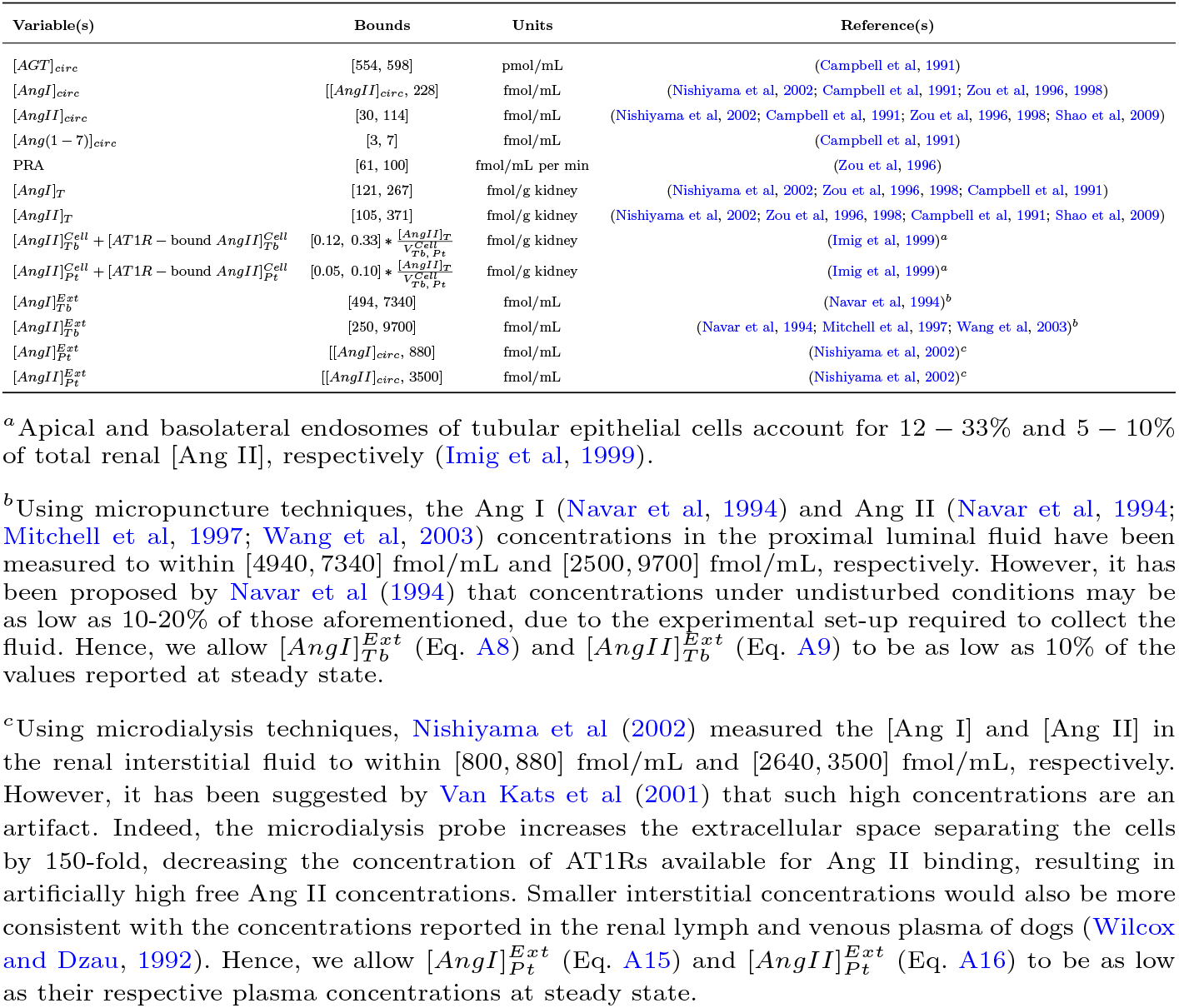
Bounds for steady state concentrations

The feedback parameters (Table V) were subsequently estimated by simulating the Ang II infusion experiments labelled *fitting* in Table AII from the initial condition *x*_0_ and minimizing the normalized sum of squared errors between the data and the simulation:

**Table V:** Feedback parameters

| Parameter | Description | Value | Unit |
| --- | --- | --- | --- |
| $k_a$ | Re-absorption rate from the subcutaneous tissue | $3.8 \times 10^{-6}$ | /min |
| $K_{circ(AGT)}$ | Strength of positive feedback on hepatic AGT production | 450 | fmol/mL/ min |
| $K_{circ(ACE)}$ | Strength of positive feedback on systemic ACE activity | 3.9 | fmol/mL per min |
| $K_{Pt}$ | Strength of positive feedback on peritubular AT1R production | 0.825 | fmol/mL per min |
| $K_{Tb}$ | Strength of positive feedback on tubular AT1R production | 0.05 | fmol/mL per min |
| $[AT1R]_{circ}^{Tot}$ | Total systemic AT1R concentration | 521500 | fmol/mL |
| $[AT1R]_{Pv}^{Tot}$ | Total blood vasculature AT1R concentration | 500 | fmol/mL |
| $B_{AT1R}$ | Strength of Ang II–AT1R feedback on renin secretion | 2.9 | – |

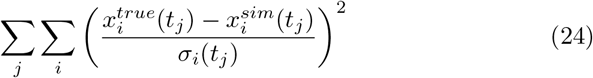

where 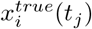 and 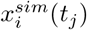 correspond to the *i*^th^ true and simulated data point collected at time point *t*_*j*_, respectively and *σ*_*i*_(*t*_*j*_) corresponds to the standard deviation of 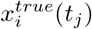.

## 3 Results

### 3.1 Model parameter identification

As outlined in section 2.4, the model parameters were fitted in a two-step process. First, we identified the baseline parameter set by minimizing Eq. 23 subject to the constraints outlined in section 2.4; see Table II. The resulting steady state concentrations *x*_0_ are shown in Table III. Then in a second-step, we simulated the Ang II infusion experiments labelled *fitting* in Table AII using the identified steady state *x*_0_ (Table III) as the initial condition. We minimized Eq. 24 to obtain the optimized feedback parameter set shown in Table V. In the cases where time series data was available, only the last time point (day 13) was used for parameter fitting. All other data points were kept for model validation (see section 3.2). Below we compare the model solutions obtained using these parameters and discuss their implications.

#### High experimental renal interstitial and luminal fluid angiotensin concentrations may be artifacts of the sample collection procedure

Using the baseline parameter set (Table II) we obtained steady state concentrations *x*_0_ (Table III) that satisfy all bounds outlined in Table IV. However, the model predicts steady state concentrations of 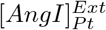 (454 fmol/mL) and 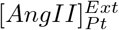 (189 fmol/mL) that are 51–57% and 5–7% of those reported by Nishiyama et al (2002), respectively. In fact, none of the parameterizations considered with peritubular interstitial angiotensin concentrations within the ranges reported by Nishiyama et al (2002) were able to satisfy the other model constraints. This discrepancy has been previously addressed by Van Kats et al (2001), who hypothesized that concentrations collected using microdialysis techniques may be artificially high due to a disturbed cell micro-environment. Moreover, the predicted concentrations of 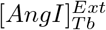 (1634 fmol/mL) and 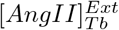 (757 fmol/mL) are 22–33% and 8–30% of those reported by Navar et al (1994), Mitchell et al (1997), and Wang et al (2003). This supports the claim made by Navar et al (1994) that in unperturbed conditions luminal fluid Ang II concentrations may be only 10–20% of those collected using micro-puncture techniques, or under 1000 fmol/mL.

#### The upregulation of hepatic AGT may influence downstream intrarenal, as opposed to systemic, RAS peptide concentrations in Ang II–induced hypertension

As shown in Fig III, the parameterized model is able to predict the changes in plasma [AGT], PRA, and [Ang I] that accompany 13 days of 40 ng/min subcutaneous Ang II infusion. Although the temporal dynamics are also predicted, only the data on day 13 was used for fitting. The remaining data was used for model validation (see section 3.2). In particular, hepatic Ang II-dependent AGT positive feedback *fb*_*circ*(*AGT*)_ (Eq. 7) was found to be required for the simulated fold-increase in plasma [AGT] to match what is reported experimentally (Zou et al, 1996) (Fig IIIa). However, given that the Michaelis constant for AGT–renin binding is so large (*K*_*M*_ = 2.8× 10^6^ fmol/mL (Gutkowska et al, 1984)), *fb*_*circ*(*AGT*)_ has an inconsequential effect on PRA (Fig IIIb) and thus, the concentration of plasma peptides downstream in the cascade. We hypothesize that the physiological importance of this feedback is instead specific to the kidney and other local RAS’ where the pro-renin receptor is expressed (Nguyen et al, 2002; Nguyen, 2006; Campbell, 2008; Wang et al, 2015): Indeed, renin binding to the pro-renin receptor (PRR) decreases the Michaelis constant for AGT binding by 85% (Nguyen et al, 2002). As a result, any substantial change in the local [AGT] will impact local renin activity, and thus the local concentrations of downstream peptides such as Ang I and Ang II. Since hepatic AGT is the primary source of AGT in the kidney (Matsusaka et al, 2012), the upregulation of hepatic (systemic) [AGT] could be one mechanism by which renal renin activity (assumed constant in our model) is maintained following Ang II infusion (Shao et al, 2009), despite the decrease in PRA that occurs.

**Fig. III:**
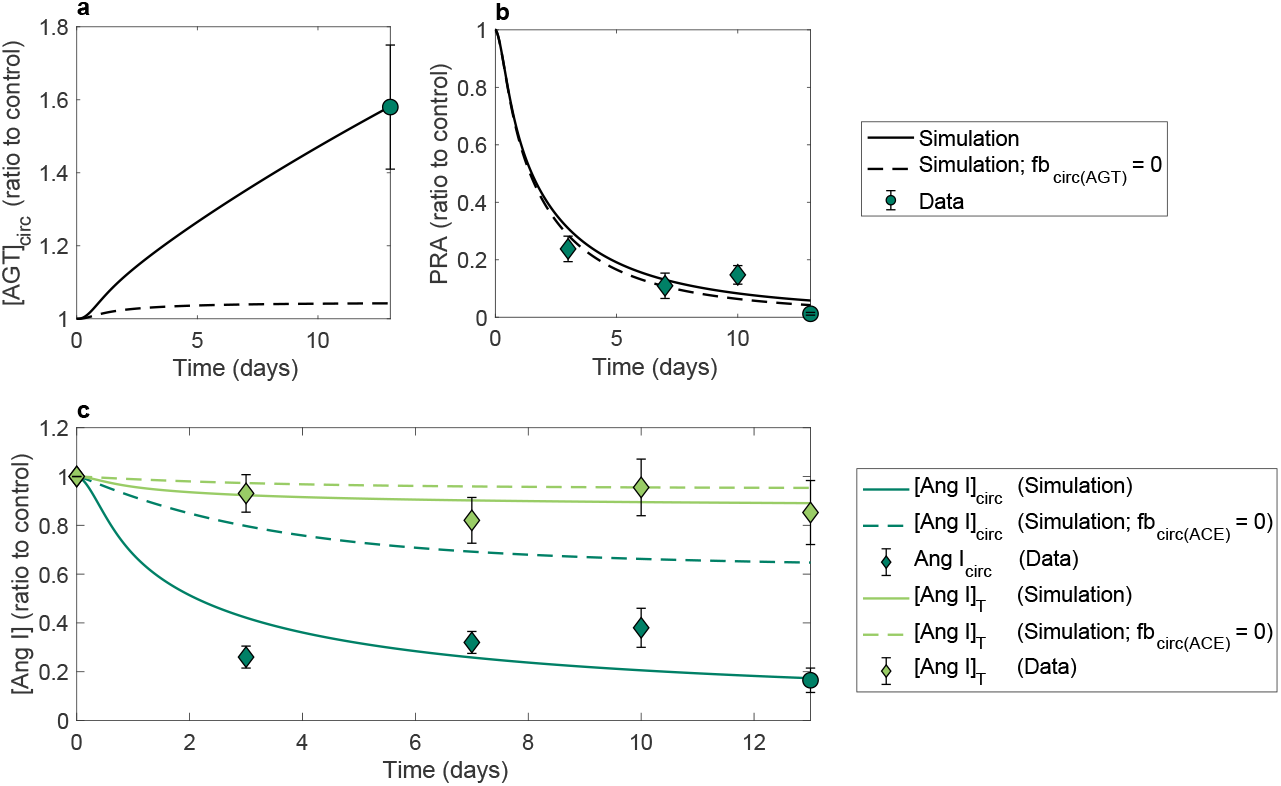
Simulated vs. experimental (a) plasma [AGT], (b) PRA, and (c) plasma (dark green) and whole kidney (light green) [Ang I] time series following 13 days of 40 ng/min SC Ang II infusion. Circular markers indicate data that was used for fitting; diamond markers indicate data used for validation. Dashed curves indicate simulations made with the specified systemic positive feedback (*fb*_*circ*(*ACE*)_ or *fb*_*circ*(*AGT*)_) removed (set to 0). Data was taken from Zou et al (1996).

#### Amplification of systemic ACE activity may be required to significantly decrease (increase) the plasma [Ang I] ([Ang II]) in Ang II–induced hypertension

As shown in Fig IIIc (dark green), the constant infusion of Ang II significantly reduces the plasma [Ang I]. Simulations suggest that this is the result of i) decreased production from AGT (inhibited renin secretion, see Eq. 16) and ii) increased conversion to Ang II (enhanced ACE activity, see Eq. 7). Of note, reduced renin activity alone was not found to be sufficient to cause the drop in plasma [Ang I] that is observed experimentally. Without enhanced ACE activity (and thus, more rapid degradation of Ang I), the plasma [Ang I] following 13 days of 40 ng/min subcutaneous infusion plateaus at 73% of its control value (Fig IIIc, dark green dashed curve), which far exceeds what the data suggests (17%). This provides additional support for the existence of the Ang II-dependent up-regulation of systemic ACE activity (*fb*_*circ*(*ACE*)_, Eq. 7).

Plasma [Ang II] dose-response curves for both classes of Ang II infusion (SC: Fig IVa and IV: Fig IVb) at three different time points (1-hour, 7 days, and 13 days) were also simulated and compared to experimental data (circular markers) to estimate the feedback parameters. As demonstrated in Fig IV, the model adequately predicts the relative increase in plasma [Ang II] that is observed experimentally. This includes the distribution of endogenously produced vs. exogenously infused plasma [Ang II] following 13 days of SC infusion at 80 ng/min (Fig IVa inset). However, the simulations appear to over-estimate the change in plasma [Ang II] caused by small SC-infused doses of Ang II. This could indicate that the positive feedback on systemic ACE activity (*fb*_*circ*(*ACE*)_, Eq. 7) depends non-linearly on the local [Ang II]. Nevertheless, linearity was assumed to avoid issues of parameter identifiability given the lack of data available at small non-vasopressor doses of Ang II. Indeed, most studies infuse large doses of Ang II in order to induce hypertension (Campbell et al, 1991).

**Fig. IV:**
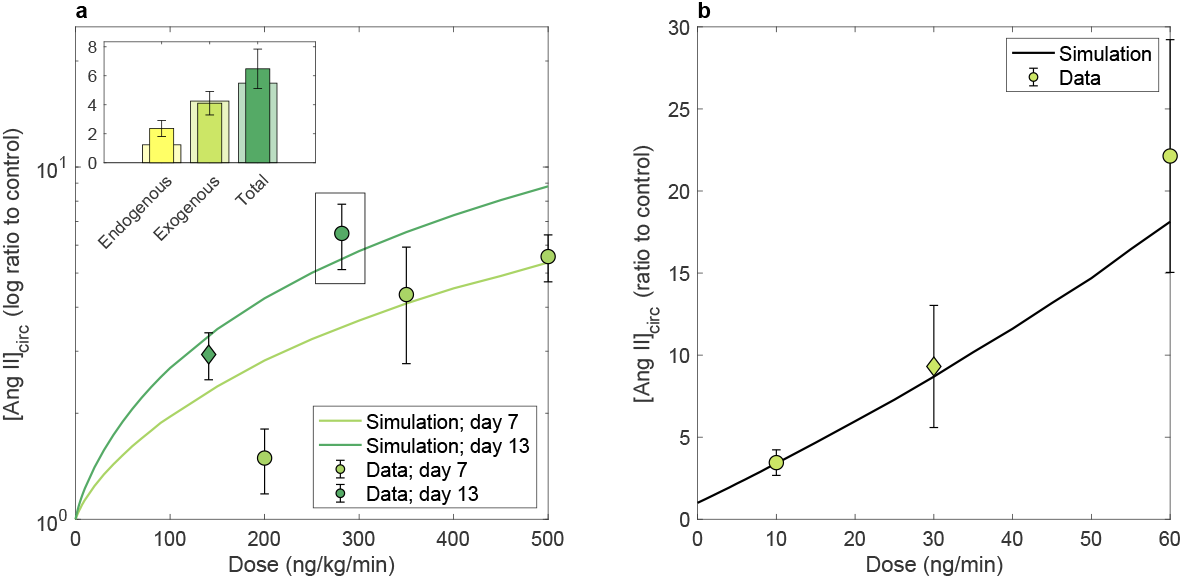
Simulated vs. experimental SC (a) and IV (b) Ang II dose–plasma [Ang II] response curves. a: Dose response following 7 (light green curve) and 13 (dark green curve) days of SC Ang II infusion. Inset: contribution of endogenous vs. exogenous Ang II to the total plasma Ang II concentration at that dose (80 ng/min ≈ 282 ng/kg/min assuming a 284 g rat). Narrow bars indicate data used for fitting; wide bars indicate the model simulation. b: Dose response following 1-hour of IV Ang II infusion. Circular (diamond) markers indicate data used for fitting (validation). All data is provided in Table AII.

#### Increased intrarenal Ang II production is likely not required for endogenous Ang II to accumulate in the kidney during Ang II infusion

The remaining data points used for parameter identification were specific to Ang II concentrations within kidney. Indeed, feedback parameters were optimized by fitting to the observed fold-change in renal endogenous and exogenous (Shao et al, 2009, 2010), interstitial (Nishiyama et al, 2003), and intracellular endosome (Zhuo et al, 2002) [Ang II] following 13 days of SC Ang II infusion at 80 ng/min. As shown in Fig Va, model estimates are within range of the experimental data in each compartment. Of note, the total renal [Ang II] after 13 days of Ang II infusion is comprised of approximately equal parts endogenous and exogenous Ang II, despite exogenous Ang II making up a larger proportion of the total plasma [Ang II] at this time (Fig IVa, inset). Interestingly, the disproportionate renal accumulation of endogenous vs. exogenous Ang II is observed despite the rates of renal Ang II production remaining constant in the model. This suggests that an explicit increase in renal Ang II production is not required for proportionally more endogenous Ang II to accumulate in the kidney in Ang II–induced hypertension.

**Fig. V:**
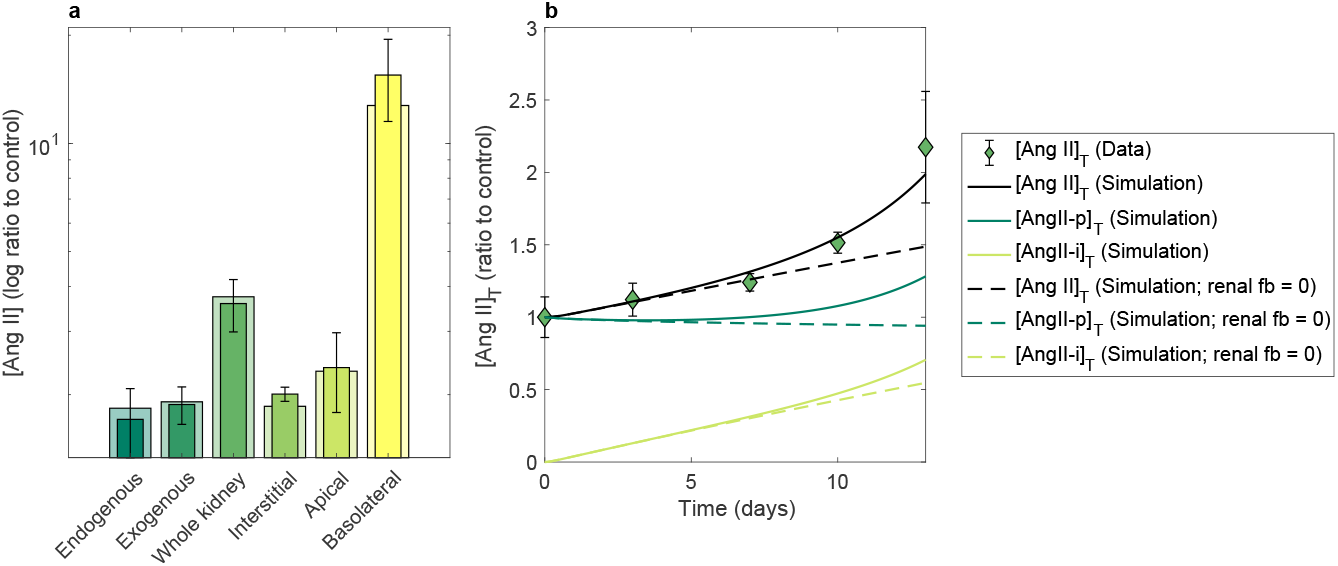
Simulated vs. experimental renal Ang II concentrations following subcutaneous Ang II infusion. a: Model fit (wide bars) to renal compartmental Ang II data (narrow bars) following 13 days of SC Ang II infusion (dose: 80 ng/min). b: Model validation (black solid curve) against whole kidney Ang II time series data following SC Ang II infusion (dose: 40 ng/min). Dark and light green curves reflect the simulated contribution of endogenous and exogenous renal Ang II to the total, respectively. The dashed curves represent concentrations simulated without renal positive feedback (*fb*_*Pt*_ = *fb*_*Tb*_ = 0 (Eq. 7)).

### 3.2 Model validation

To validate the model, we simulate the Ang II infusion experiment carried out by Zou et al (1996) and compare the model predictions to data. In particular, Zou et al (1996) subcutaneously infused Ang II at 40 ng/min for 13 days and observed a rapid, slight decrease in the total renal [Ang I], as well as slow rise in the total renal [Ang II]. The model is able to closely capture these results, as shown in Figs IIIc (light green curve) and Vb (black curve), respectively. Model simulations also predict the behaviour of plasma peptides, such as: the fold increase in plasma [Ang II] observed at the end of the experiment (day 13) and the fold-decrease in PRA and plasma [Ang I] observed 3, 7, and 10 days into the study. Results are shown in Figs IVa (diamond marker), IIIb, and IIIc (dark green), respectively.

In the next section, we examine the experimental results presented by Zou et al (1996) in more detail and use model simulations to offer insight into the underlying mechanisms.

### 3.3 Model predictions

#### Renal [Ang I] decreases throughout the development Ang II–induced hypertension because of a reduction in plasma [Ang I], not renal renin activity

As shown in Fig IIIc, renal [Ang I] decreases significantly less than plasma [Ang I] over the course of the low-dose Ang II infusion. Model simulations can be used to explain this behaviour. Indeed, the large decrease in plasma [Ang I] (Fig IIIc, dark green curves) that accompanies Ang II infusion was found to be sufficient to cause the small decrease in renal [Ang I] (Fig IIIc, light green curves) that is observed experimentally; no decrease in local endogenous production (assumed constant in the model) was required. This is consistent with the experimental observation that renal renin activity is conserved during Ang II infusion (Shao et al, 2009). Possible mechanisms contributing to the maintenance of kidney renin concentration following Ang II infusion are i) increased renin production in the collecting duct (Prieto-Carrasquero et al, 2004; Navar et al, 2011; Gonzalez and Prieto, 2015; Gonzalez et al, 2011) and ii) local PRR expression in conjunction with increased [AGT] from elevated hepatic (see section 3.1) (Zou et al, 1996; Matsusaka et al, 2012) and proximal tubule (Navar et al, 2003, 2011; Kobori et al, 2001, 2004) AGT production.

#### AT1R–mediated uptake of Ang II is the primary mechanism by which Ang II accumulates in the kidney in Ang II–induced hypertension

By simulating the Ang II infusion experiment carried out by Zou et al (1996) in the absence and presence Ang II–dependent positive feedback, we can gain insight into its role in the development of Ang II–induced hypertension. Indeed, the model predicts that Ang II-dependent positive feedback on AT1R expression is particularly important in the kidney during the second week of low-dose (40 ng/min) SC Ang II infusion: During the first week, exogenously-infused Ang II (Fig Vb, light green curves) alone accounts for the majority of the increase in total renal [Ang II], even with all renal positive feedback removed (Fig Vb, dashed curves). This is no longer true in the second week of infusion, where up-regulated AT1R expression is required for a sufficient concentration of endogenous Ang II (Fig Vb, dark green curves) to accumulate in the kidney. Both AT1R-mediated uptake of Ang II and increased local production of Ang II have been suggested as mechanisms driving increased renal [Ang II] in Ang II-dependent hypertension (Kobori et al, 2007; Navar et al, 2011; Shao et al, 2009, 2010; Van Kats et al, 2001; Zhuo et al, 2002; Kobori et al, 2001; Ferrão et al, 2014; Gonzalez-Villalobos et al, 2008). Our results suggest that AT1R-mediated uptake of Ang II is the primary mechanism by which Ang II accumulates in the kidney and that increased local expression of AT1Rs, not Ang II, is required.

Given that the renal mechanisms that affect blood pressure are compartment specific, local changes to the distribution of renal angiotensin peptides may lead to blood pressure dis-regulation. Therefore, in the next section we investigate the distributional changes in renal [Ang II] that accompany Ang II infusion to gain insight into the development of Ang II–induced hypertension.

#### The accumulation of Ang II in tubular epithelial cells is likely crucial to the onset of Ang II–induced hypertension

Fig VI illustrates the predicted temporal change in the renal distribution of [Ang II] during low-dose (40 ng/min) Ang II infusion. In a normotensive rat, the tubular compartment makes up the majority of the total renal [Ang II] (Fig VIa, time 0), with the highest concentration of peptides in the luminal fluid (Fig VIb, time 0). As shown in Fig VI, this does not change significantly over the first 5 days of low-dose SC Ang II infusion. However, at this point the exogenously infused Ang II has sufficiently increased the concentration of apical and basolateral-bound AT1R–Ang II complexes 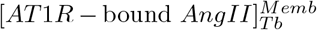 and 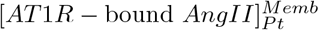 over their steady states to activate the positive feedback on AT1R expression (*fb*_*Tb*_ > 0 and *fb*_*Pt*_ > 0 (Eq. 7)). As a result, a feed-forward loop is initiated, whereby increased AT1R expression leads to more AT1R–Ang II binding (Fig VIc) which results in more AT1R expression, and so on. Ultimately, more AT1R–Ang II complexes become internalized through both the apical and basolateral membranes of tubular epithelial cells, causing Ang II to accumulate in this intracellular compartment (Fig VId). In particular, since the strength of the positive feedback on basolateral AT1R expression *K*_*Pt*_ is greater than that of apical AT1R expression *K*_*Tb*_ (Eq. 7), a greater proportion of Ang II gets accumulated via the basolateral membrane i.e. within the peritubular compartment. Interestingly, the time at which Ang II starts accumulating in the peritubular and tubular intracellular compartments of the model, i.e. in tubular epithelial cells, (day 6) coincides exactly with when the rats in Zou et al (1996)’s study started exhibiting a detectable increase in systolic blood pressure. This indicates that the accumulation of Ang II in tubular epithelial cells may play a key role in the development of Ang II–induced hypertension, likely through the stimulation of sodium reabsorption (Hall, 1986; Schuster et al, 1984; Harris and Young, 1977; Schelling and Linas, 1994).

**Fig. VI:**
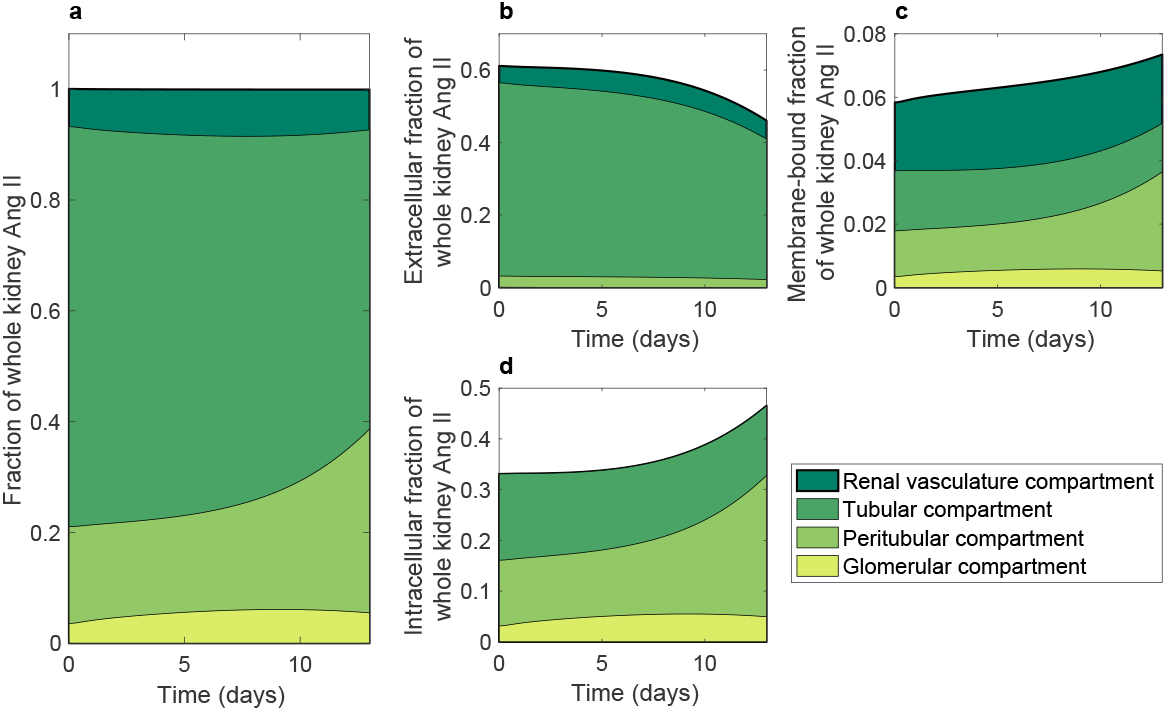
Temporal change in the renal distribution of [Ang II] throughout 13 days of SC Ang II infusion at 40 ng/min. Panels depict the relative contribution of each compartment to the: (a) total renal [Ang II], (b) extracellular fraction of renal [Ang II], (c) membrane-bound fraction of renal [Ang II], and (d) intracellular fraction of renal [Ang II].

To determine the robustness of the results presented in sections 3.1, 3.2, and 3.3, we perform a local parametric sensitivity analysis.

### 3.4 Local sensitivity analysis

We examine the percent change in the model variables at steady state and after 13 days of simulating the Ang II infusion experiment performed by Zou et al (1996) when each parameter is increased by 10%.

As shown in Fig VII, the steady state is most (least) sensitive to changes in the baseline parameters specific to the tubular (peritubular) compartment, in particular, the rates of endogenous Ang I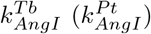 and Ang II 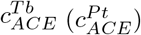 production in the luminal (interstitial) fluid. The model is also very sensitive to the proportion of Ang I and Ang II fluxed across the tubular epithelium *S*_*Tb*_. Indeed, the majority of the renal variables (apart from those in the upstream glomerular compartment) are altered by this parameter. Finally, the behaviour of the systemic, glomerular, and renal (blood) vasculature free [Ang I] and [Ang II] steady states appear to be correlated. Specifically, since systemic angiotensin is the main source of glomerular and renal (blood) vasculature angiotensin, the variables in these compartments are most affected by changes to parameters relating to systemic peptide production (*k*_*AGT*_, *v*_*max*_, and *c*_*chym*_). Nevertheless, a 10% change in each parameter never elicits more than a 10% change any model variable and as a result, the model’s steady state remains within the bounds presented in Table IV in all cases considered.

**Fig. VII:**
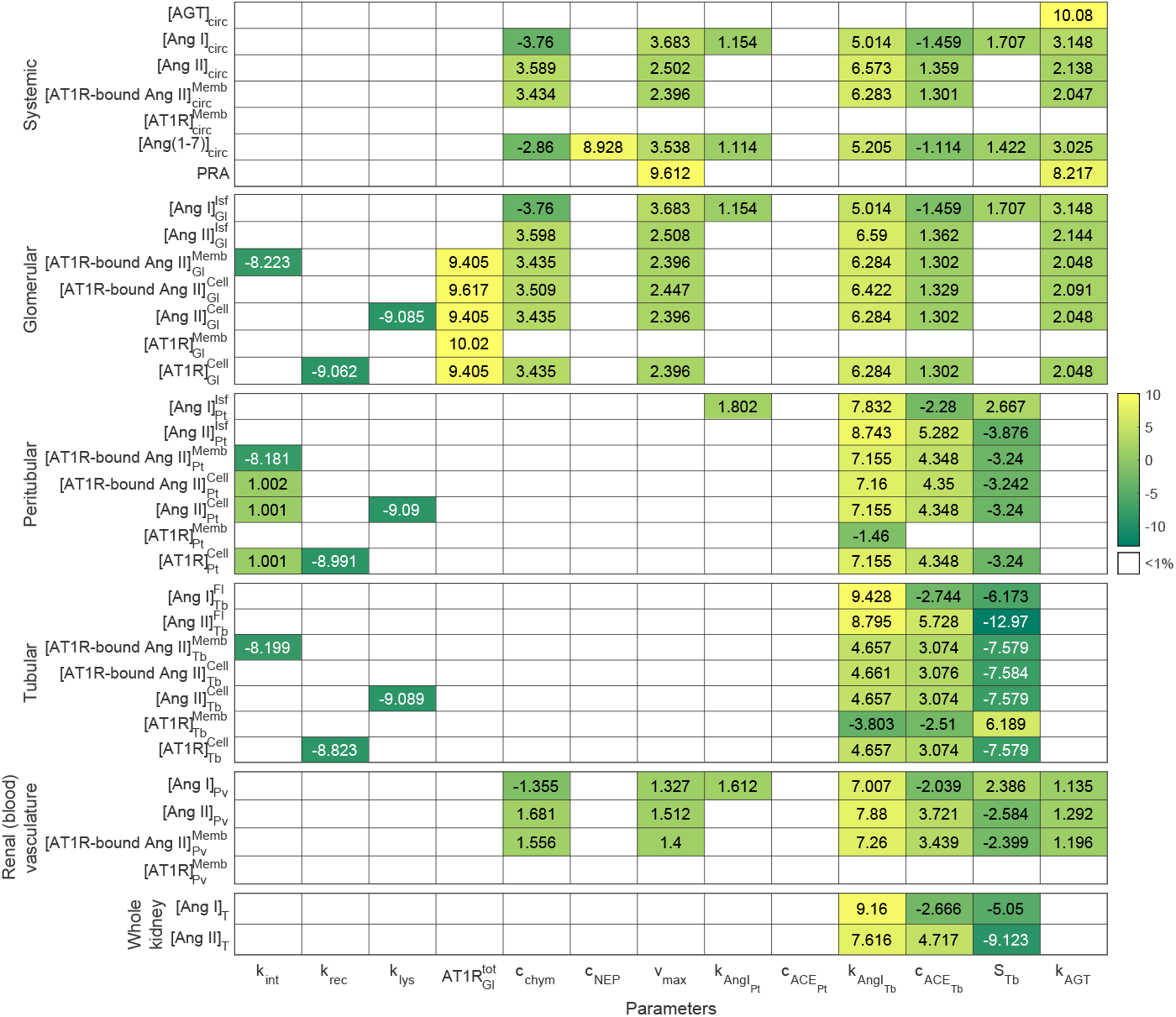
Percent change in the model steady state caused by a 10% increase in each baseline parameter value.

As shown in Fig VIII, the model predictions following 13 days of SC 40 ng/min Ang II infusion are most sensitive to changes in the same baseline parameters that the steady state was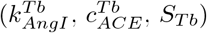. Moreover, parameter changes that increase the rate of endogenous Ang II production at baseline (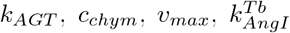, and 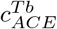) often decrease the concentration of tubular epithelial cell-associated Ang II. This is because the same amount of exogenous Ang II is entering these compartments, but there is more endogenous AT1R–bound Ang II at baseline. Hence, any exogenous AT1R–Ang II binding elicits a smaller fold-increase in the AT1R–bound Ang II concentration which blunts the positive feedback on AT1R expression *fb*_*Tb*_ and *fb*_*Pt*_ (Eq. 7) and leads to less cell-associated Ang II.

**Fig. VIII:**
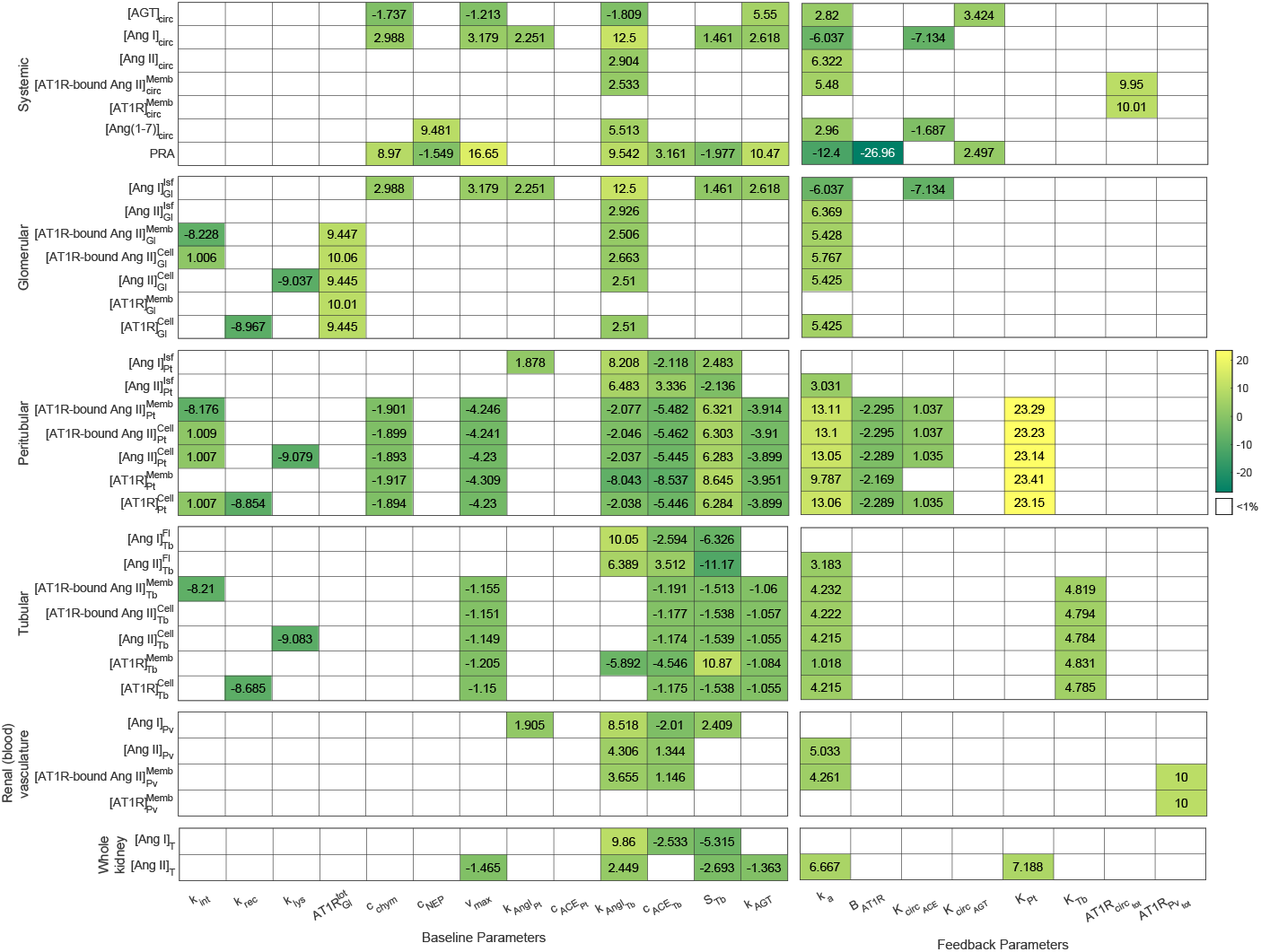
Percent change of the model predictions following 13 days of 40 ng/min SC Ang II infusion to a 10% increase in each baseline and feedback parameter value.

In terms of the feedback parameters, an increase in the rate of Ang II reabsorption from the subcutaneous tissue *k*_*a*_ has the most significant impact on the model predictions following Ang II infusion (Fig VIII). Indeed, the faster the exogenous Ang II is reabsorbed, the sooner all positive feedback gets activated allowing for greater accumulation of Ang II in the kidney and greater endogenous production in the plasma. The only other feedback parameter that notably affects the accumulation of Ang II in the kidney during infusion is the strength of the positive feedback on AT1R expression along the basolateral membrane of tubular epithelial cells *K*_*Pt*_. Indeed, a 10% increase in this parameter elicits a 23% increase in cell-associated Ang II in the peritubular compartment. As a result, 7% more Ang II accumulates in the kidney during the Ang II infusion. A similar effect is observed when the strength of the AT1R expression feedback along the apical membrane *K*_*Tb*_ is increased. However, given its smaller original value, a 10% increase in this parameter elicits only a 4% change in cell-associated Ang II in the tubular compartment. This isn’t enough to significant alter the whole kidney [Ang II].

## 4 Discussion

The primary goal of this paper was to gain insight into the role of the intrarenal RAS in the development of hypertension. To accomplish this, a mathematical model of the intrarenal RAS was created, parameterized, validated, and used to simulate Ang II infusion experiments. In particular, the low dose (40 ng/min), 13 day subcutaneous Ang II infusion experiment carried out by Zou et al (1996) was replicated and the results were examined in detail.

AT1R–mediated uptake of circulating Ang II and enhanced endogenous Ang II generation have both been suggested as key mechanisms contributing to enhanced renal accumulation of Ang II during the development of hypertension (Kobori et al, 2007; Navar et al, 2011; Shao et al, 2009, 2010; Van Kats et al, 2001; Zhuo et al, 2002; Kobori et al, 2001; Ferrão et al, 2014; Gonzalez-Villalobos et al, 2008). The model simulations suggest that AT1R–mediated uptake of Ang II into tubular epithelial cells is the primary mechanism by which Ang II accumulates in the kidney during low-dose Ang II infusion, and that enhanced local Ang II production is not required (see below). This is consistent with the findings of Zhuo et al (2002), who found that blocking AT1R–Ang II binding via angiotensin-receptor blocker (ARB) administration not only prevents the accumulation of Ang II in renal intracellular endosomes, but in the entire kidney itself. We also determined that this effect requires an Ang II–mediated increase in AT1R expression, mainly along the basolateral membrane of these cells. This result is also corroborated by Zhuo et al (2002). Indeed, Zhuo et al (2002) found that increased AT1R expression in renal endosomes is, at least in part, responsible for the observed increase in intracellular trafficking of Ang II into renal intracellular endosomes following Ang II infusion. An Ang II–dependent increase in proximal tubule AT1R expression has also been reported by other sources (Zhuo et al, 1999; Cheng et al, 1995), though to our knowledge, its functional role had yet to be discussed until now. Our results suggest that this feedback is likely crucial to the development of hypertension because it permits the intracellular accumulation of Ang II in tubular epithelial cells (see below).

Notably, Ang II was able to accumulate in the simulated kidney without the need to explicitly increase the local endogenous production of Ang II. In fact, more endogenous than exogenous Ang II accumulated in tubular epithelial cells once AT1Rs were up-regulated because the predicted basal rate of endogenous production of Ang II in the kidney exceeded the rate of exogenous Ang II entry into the kidney. An explicit increase to the rate of endogenous Ang II production was not required for this to be true. In particular, a constant renal renin activity was sufficient because it ensured that the renal [Ang I] and thus the local endogenous production of Ang II decreased only slightly during the infusion, despite the significant decrease in plasma [Ang I]. It is thus likely that the Ang II–dependent AGT amplification mechanism that has been reported in the proximal tubule (Navar et al, 2003, 2011; Kobori et al, 2001; Gonzalez-Villalobos et al, 2008; Kobori et al, 2004; Schunkert et al, 1992) primarily contributes to the maintenance of renal renin activity (Shao et al, 2009). It does not necessarily result in increased endogenous Ang II production above baseline. An elevated renal AGT concentration is able to influence renal renin activity because the kidney expresses pro-renin receptors that reduce the Michaelis constant for renin–AGT binding (Nguyen et al, 2002; Nguyen, 2006; Campbell, 2008; Wang et al, 2015). This is not the case in the plasma, where we showed that hepatic AGT amplification (Li and Brasier, 1996; Schunkert et al, 1992; Nakamura et al, 1990) had an inconsequential effect on PRA and thus, the systemic RAS. We thus hypothesize that this systemic positive feedback mainly influences the intrarenal RAS, since hepatic AGT has been reported as the primary source of AGT in the kidney (Matsusaka et al, 2012).

In addition to the hepatic AGT positive feedback (Li and Brasier, 1996; Schunkert et al, 1992; Nakamura et al, 1990), model simulations suggest that systemic ACE activity is likely also up-regulated by Ang II. Firstly, while sufficient endogenous Ang II was able to accumulate in the kidney without increasing local Ang II production, this was no longer the case in the plasma. Indeed, without enhanced systemic Ang II production, the experimentally measured endogenous concentration of plasma Ang II (Shao et al, 2009, 2010) was under-predicted by the model. Secondly, the reduction in PRA resulting from the Ang II-dependent feedback on renin secretion from the juxtamedullary apparatus was not sufficient to cause the drop in plasma [Ang I] that was observed experimentally by Zou et al (1996). Indeed, without enhanced ACE activity (and thus, more rapid degradation of Ang I), the plasma [Ang I] far exceeded what the data suggested. While similar feedback has been documented in the kidney (Sadjadi et al, 2005; Koka et al, 2008), future experiments are required to confirm its existence in the systemic circulation.

Finally, we found that the simulated onset of Ang II accumulation in tubular epithelial cells coincided exactly with the experimental inception of hypertension on day 6 of the Ang II infusion experiment performed by Zou et al (1996). The model thus suggests that the delayed rise in blood pressure is the result of the time it takes for the positive feedback on renal AT1R expression to be sufficiently activated by exogenous Ang II and consequently, for Ang II to begin accumulating within tubular epithelial cells. Hence, it is likely that the stimulation of sodium reabsorption (Hall, 1986; Harris and Young, 1977; Schuster et al, 1984; Schelling and Linas, 1994) following the association of Ang II with these cells plays a crucial role in the development of hypertension. As discussed below, this hypothesis can be further explored in future work by coupling our intrarenal RAS model to Ahmed and Layton (2020)’s whole-body model of blood pressure regulation that considers cardiovascular function, renal hemodynamics, renal sodium and fluid handling, the renal sympathetic nervous system, and the connections between these systems.

### Model limitations and future extensions

The main limitation of this model is that it considers the intrarenal and systemic RAS in isolation, when in reality these systems influence and are influenced by many other physiological processes. For example, an increase in renal Ang II is known to affect renal hemodynamic function by increasing afferent and efferent arteriole resistance (Ahmed and Layton, 2020; Yang et al, 2011; Denton et al, 2000). This results in an increased filtration fraction (lower renal blood flow, sustained glomerular filtration rate) (Toke and Meyer, 2001; Denton et al, 2000), which will in turn affect the intrarenal distribution of Ang II. This cascade of events is not captured by the present model, which assumes that all renal hemodynamic parameters are known *a priori* and as such are unaffected by Ang II infusion. In future work, the model can be extended to consider all renal hemodynamic flow rates and volumes as variables instead of parameters. In this way, the interplay between Ang II and renal hemodynamics can be incorporated and investigated in detail.

A key consequence of the model’s limited scope is that the downstream effects of altered systemic and renal RAS peptide concentrations are not modelled explicitly. We are thus forced to make assumptions about blood pressure based on prior knowledge of the processes affected by local Ang II concentrations (e.g. sodium reabsorption), without modelling mean arterial pressure directly. This limitation can be overcome in future work by incorporating the present intrarenal RAS model into a whole-body blood pressure regulation model, such as the one presented by Ahmed and Layton (2020). Using the resulting more comprehensive model, the effect of Ang II accumulation in tubular epithelial cells on mean arterial pressure can be studied directly. Coupling the models also resolves the aforementioned disconnect between altered intrarenal Ang II concentrations and altered renal hemodynamics, since the model presented by Ahmed and Layton (2020) already takes the effect of Ang II on renal vascular resistance into consideration.

In general, a whole-body blood pressure regulation model that includes the intrarenal RAS would be an excellent tool to study hypertension. With only the systemic RAS represented in Ahmed and Layton (2020)’s model, all RAS-mediated blood pressure regulation processes in the kidney (e.g., sodium reabsorption, renin secretion, afferent and efferent arteriole constriction) are assumed to be regulated directly by plasma Ang II. In reality, these processes depend on local intrarenal Ang II. This assumption becomes particularly problematic in an investigation of hypertension, given that the behaviour of intrarenal Ang II differs significantly from plasma Ang II following Ang II infusion and in other experimental models of hypertension (Zou et al, 1996; Cervenka et al, 1999; Wu et al, 2014; Takenaka et al, 1990). Therefore, incorporating the intrarenal RAS model will not only improve Ahmed and Layton (2020)’s model, but facilitate a great opportunity to study hypertension.

The effects of anti–hypertensive therapies on the intrarenal RAS can also be studied in the future by creating pharmacokinetic models of various drugs of interest and coupling them to the intrarenal RAS model presented here. Indeed, models can be created that consider drug dynamics across four compartments: the gastrointestinal tract, the systemic circulation, the poorly perfused organs, and the kidney. Two classes of drugs to consider are angiotensin-receptor blockers (ARBs) and angiotensin converting enzyme inhibitors (ACEi). Coupling to the intrarenal RAS model can be achieved through systemic and renal ARB–AT1R binding and ACEi–based inhibition of ACE activity in compartment 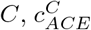. We hypothesize that ARBs and ACEi both prevent Ang II–induced hypertension, at least in part, by blocking the AT1R–mediated accumulation of Ang II in tubular epithelial cells. The drugs achieve this via different mechanisms: ACEi reduce the endogenous production of Ang II in the kidney and thus, the pool of Ang II that can be internalized. ARBs prevent Ang II–AT1R binding, and thus the internalization of Ang II altogether. We expect ARBs to be more effective than ACEi at preventing hypertension induced by Ang II infusion. This is because ACEi do not prevent the accumulation of exogenous Ang II in tubular epithelial cells following Ang II infusion, but ARBs do.

By fine-tuning its excretory function, the kidney plays a predominant role in blood pressure regulation (Guyton, 1991). Indeed, transplantation studies have demonstrated that hypertension follows the kidney (Guidi et al, 1996). As such, the mechanisms by which intrarenal RAS regulate blood pressure can be examined by coupling the present model to a kidney function model at the transporter level, such as those by Layton and Layton (2019), Layton et al (2017), Edwards et al (2014), Edwards and Layton (2014), and Chen et al (2011). The coupling can be formulated based on the known connections between Ang II, AT1Rs, and ion transport (Burns and Li, 2003; Valles et al, 2005).

Finally, the current intrarenal RAS model does not consider sex differences, even though it has been long established that the steady state concentrations of most systemic RAS peptides differ between males and females (Pendergrass et al, 2008), and each sex responds differently to antihypertensive therapies that target the RAS (Leete et al, 2018). To account for and explain these disparities, Leete et al (2018) parameterized separate computational models of the systemic RAS for male and female rats. These models were later integrated into the whole body blood pressure regulation models by Karaaslan et al (2005) and Hallow et al (2014) to create sex-specific models of blood pressure regulation in both humans (Leete and Layton, 2019) and rats (Ahmed and Layton, 2020). In future work, our intrarenal RAS model can be extended to account for the known sex differences that exist both at the systemic and intrarenal level using the experimental measurements collected by Pendergrass et al (2008). Furthermore, the model can be used to help infer sex-differences that exist at the compartmental level (tubular fluid, interstitium, vasculature, etc.), based on the whole-tissue peptide concentrations that are available (Pendergrass et al, 2008). The sex-specific intrarenal RAS models could also later be coupled to a sex-specific blood pressure regulation model (Ahmed and Layton, 2020) or to a sex-specific epithelial solute transport models (Li et al, 2018a; Hu et al, 2019, 2020, 2021) to study whether the roles of the intrarenal RAS in hypertension and renal function differ between the sexes.

## Conclusion

In this work, we have discussed the role of the intrarenal RAS in the pathogenesis of hypertension induced by Ang II infusion. However, Ang II infusion is an experimental model which relies on the constant administration of non physiological doses of Ang II (Yang and Xu, 2017). As a result, it has a limited scope in terms of its relevance to clinical hypertension. Nevertheless, the conclusions drawn from this work may have broader implications to other forms of hypertension that are associated with an overactive RAS. Indeed, the decoupling of the systemic and intrarenal RAS has been observed in many other experimental models of hypertension that better represent clinical hypertension, such as: two-kidney, one-clip Goldblatt hypertension (Cervenka et al, 1999), salt-sensitive rats (Wu et al, 2014), and spontaneously hypertensive rats (Takenaka et al, 1990). Hence, it is expected that the systemic and intrarenal RAS become decoupled in cases of clinical hypertension that are associated with an over-active RAS. Furthermore, the mechanisms mediating this effect are likely the same as those that contribute to Ang II–induced hypertension. These hypotheses can be further explored in future work, by using the presented model to study the explicit functional role of the intrarenal RAS in other experimental models of hypertension that do no involve the exogenous infusion of Ang II.

## Statements and declarations

### Code availability

https://github.com/Layton-Lab/intrarenalRAS.git

## Appendix A

### A.1 Intrarenal RAS model equations

Below we summarize model equations for each compartment.

#### A.1.1 Glomerular compartment

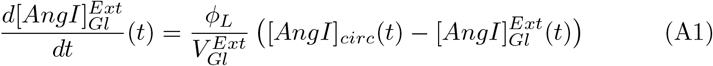

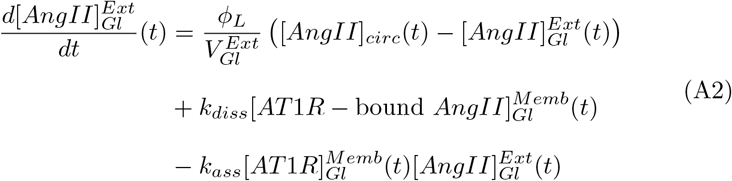

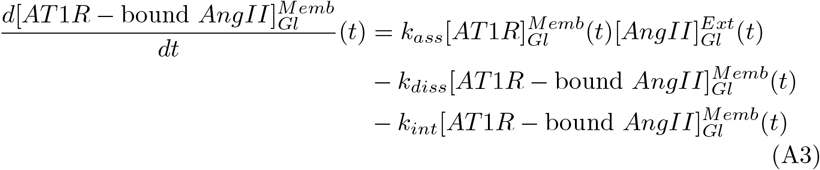

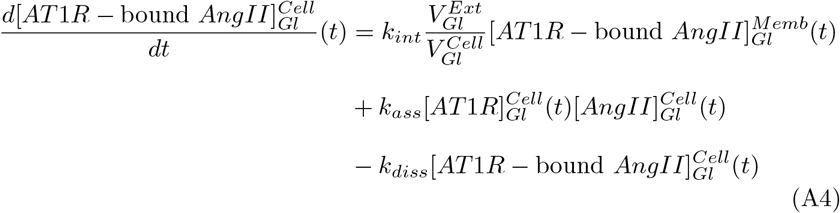

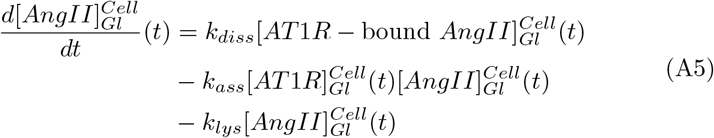

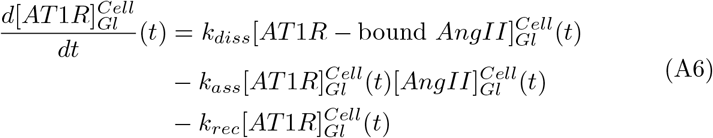

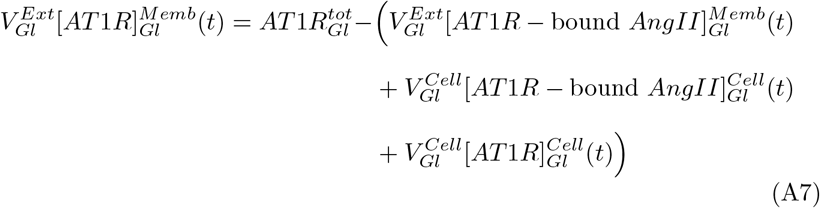

#### A.1.2 Tubular compartment

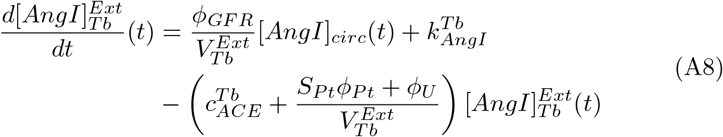

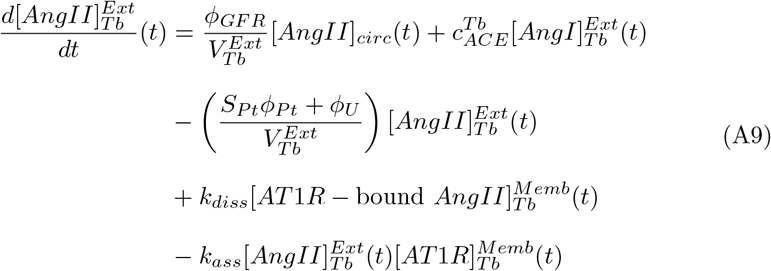

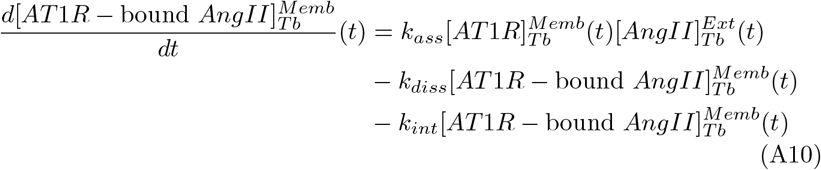

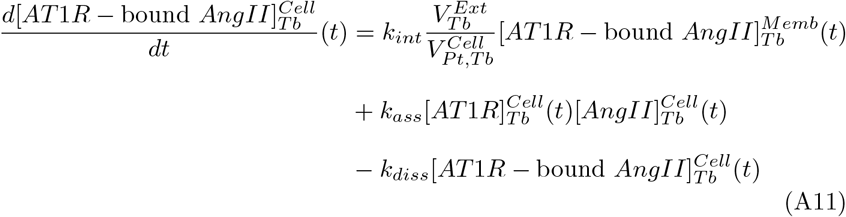

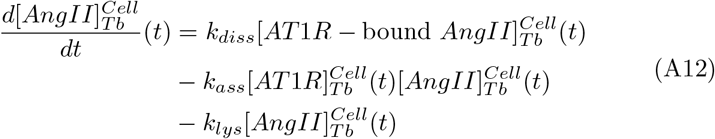

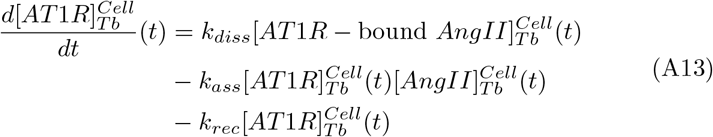

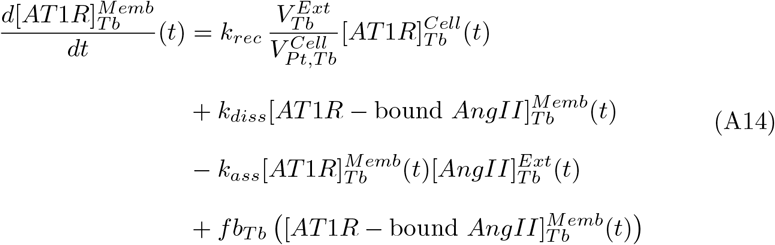

#### A.1.3 Peritubular compartment

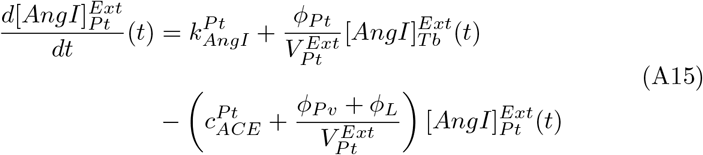

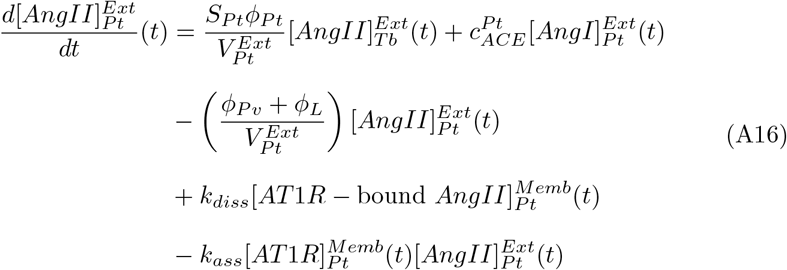

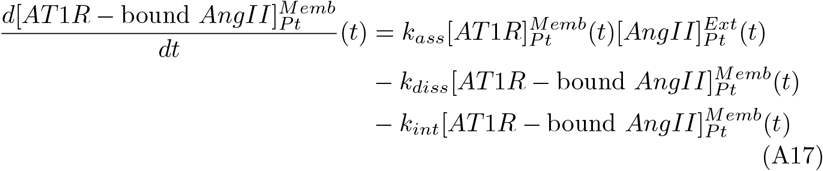

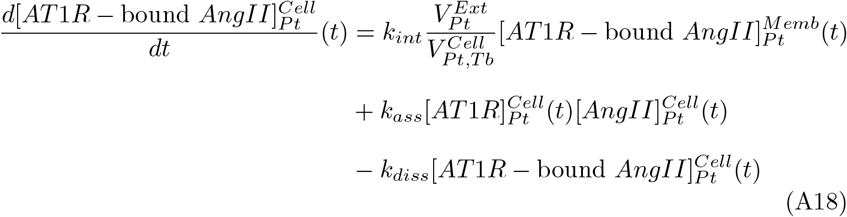

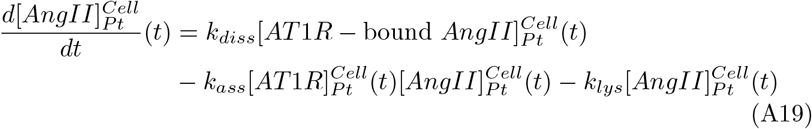

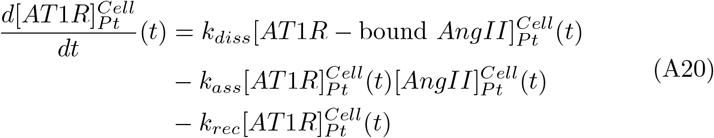

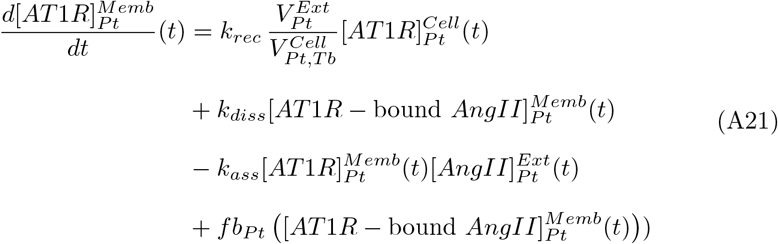

#### A.1.4 Blood vasculature compartment

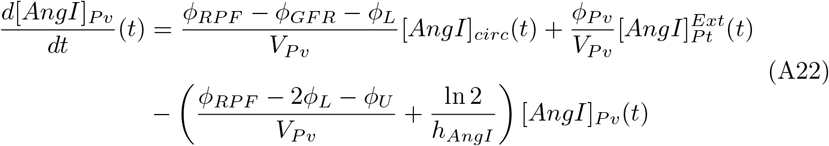

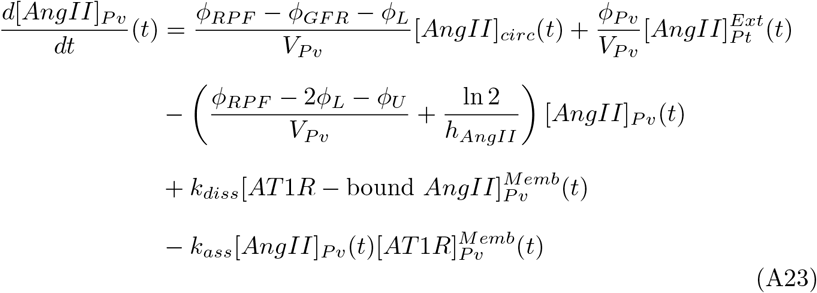

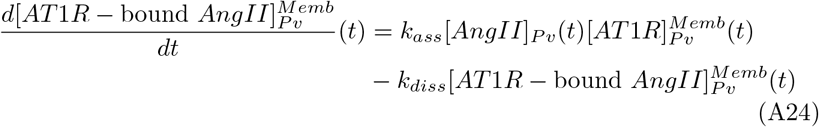

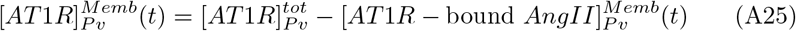

### A.2 Baseline parameter derivations

The model parameters listed in Table AI were derived from the literature in the following way:

1. **General parameters:** Many whole-body parameters are based on the 284 g male Sprague Dawley rat described by Munger and Baylis (1988). Indeed, given a kidney-to-body weight ratio of 5.23 (Zou et al, 1996), we assume a kidney weight *W*_*K*_ of 1.49g. Based on the body weight-to-blood volume relationship from Lee and Blaufox (1985), we assume a circulating blood volume of 17.8mL. This corresponds to a circulating plasma volume *V*_*circ*_ of 10.3 mL, given a hematocrit of 0.42 (Munger and Baylis, 1988).
2. **Renal volume parameters:** We extract the cortical volume shares of the luminal fluid (0.16), tubular epithelial cells (0.46), and the interstitial tissue including peritubular capillaries (0.30) from Hegedüs et al (1978). Moreover, we know the cortical volume share of the peritubular interstitium (0.037) and entire peritubular interstitium including interstitial cells (0.07) from Lemley and Kriz (1991), allowing us to compute that of the peritubular capillaries (renal blood vasculature) alone (0.30 − 0.07 = 0.23). Since the kidney is 70% cortex by volume (Baldelomar et al, 2018), and the ratio of kidney volume-to-kidney weight is 0.912 mL/g (extracted from Fig 3F in Baldelomar et al (2018)), we obtain:

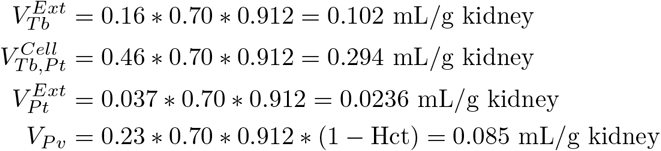

The volumes of the glomerular interstitium and intracellular compartments are computed based on the known volume fractions in mice (Katz et al, 2002). Indeed, with a mesangial matrix and mesangial cell-to-glomerulus fractional volume of 0.09, a glomerulus-to-whole kidney fraction volume of 0.023, and a kidney volume-to-kidney weight ratio of 0.912 (Baldelomar et al, 2018), we obtain:

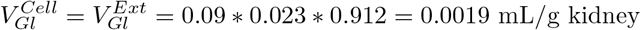
3. **Renal hemodynamic parameters:** Renal plasma flow *ϕ*_*RPF*_ and glomerular filtration rate *ϕ*_*GFR*_ are taken directly from Munger and Baylis (1988) and scaled by *W*_*K*_ to obtain the appropriate units of mL/g kidney. Moreover, we assume that renal lymphatic flow i) accounts for 2% of total fluid reabsorption from the kidney (*ϕ*_*L*_ = 0.02*ϕ*_*Pt*_) (Sugarman et al, 1942) and ii) that it is the same as urine flow (*ϕ*_*U*_ = *ϕ*_*L*_) (Russell et al, 2019). In this way, all remaining hemodynamic parameters can be computed from the known glomerular filtration rate (Munger and Baylis, 1988), since *ϕ*_*L*_ = *ϕ*_*U*_ = (0.02*/*1.02)*ϕ*_*GFR*_, *ϕ*_*Pt*_ = *ϕ*_*GFR*_−*ϕ*_*U*_, and *ϕ*_*Pv*_ = *ϕ*_*GFR*_−*ϕ*_*L*_−*ϕ*_*U*_.

### A.3 Ang II infusion data

The data in Table AII was collected from a wide variety of Ang II-infusion studies, and used to fit the feedback parameters and validate the model.

**Table AI:**
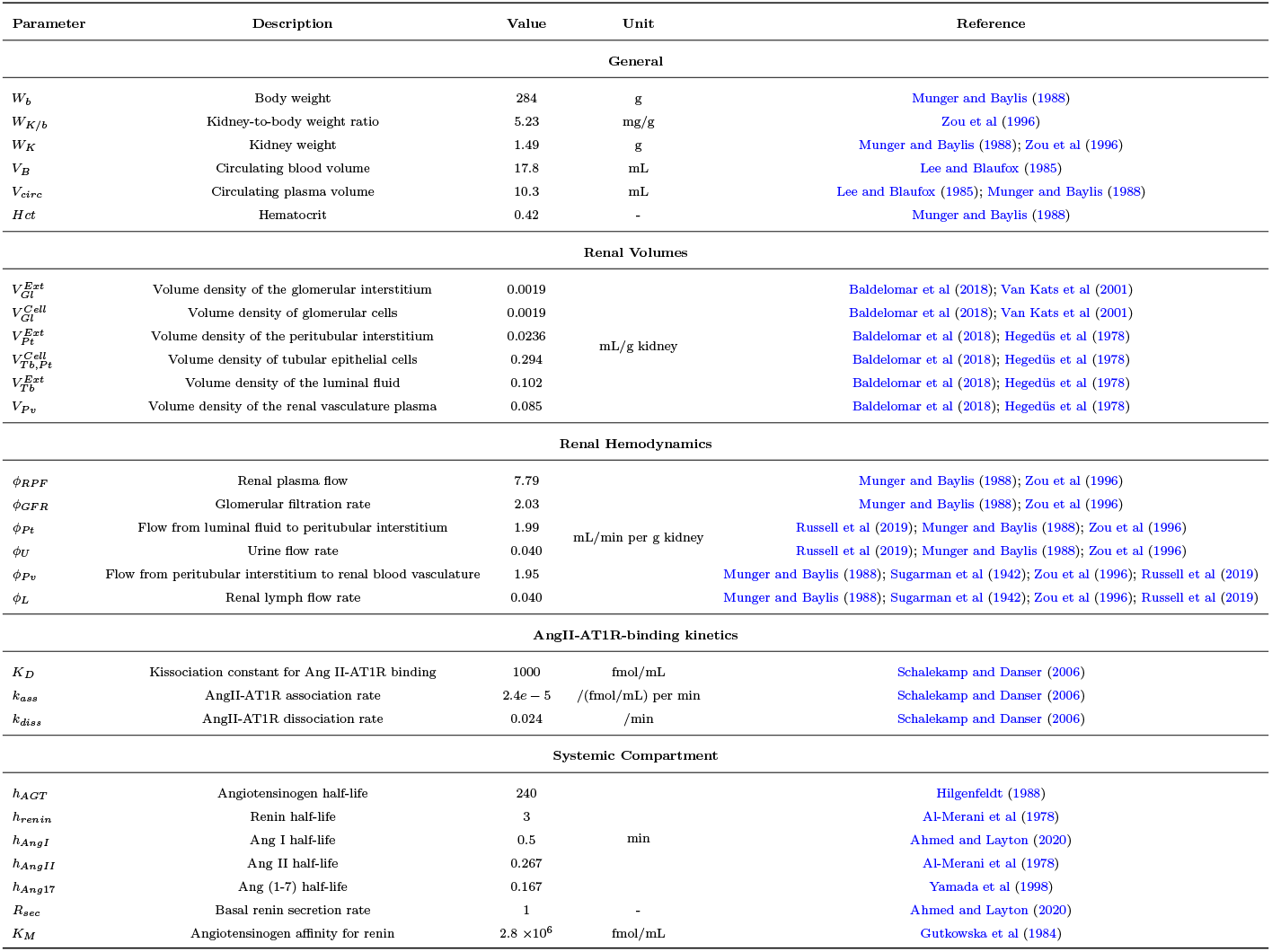
Baseline parameters derived from the literature

---

**Table AII:**
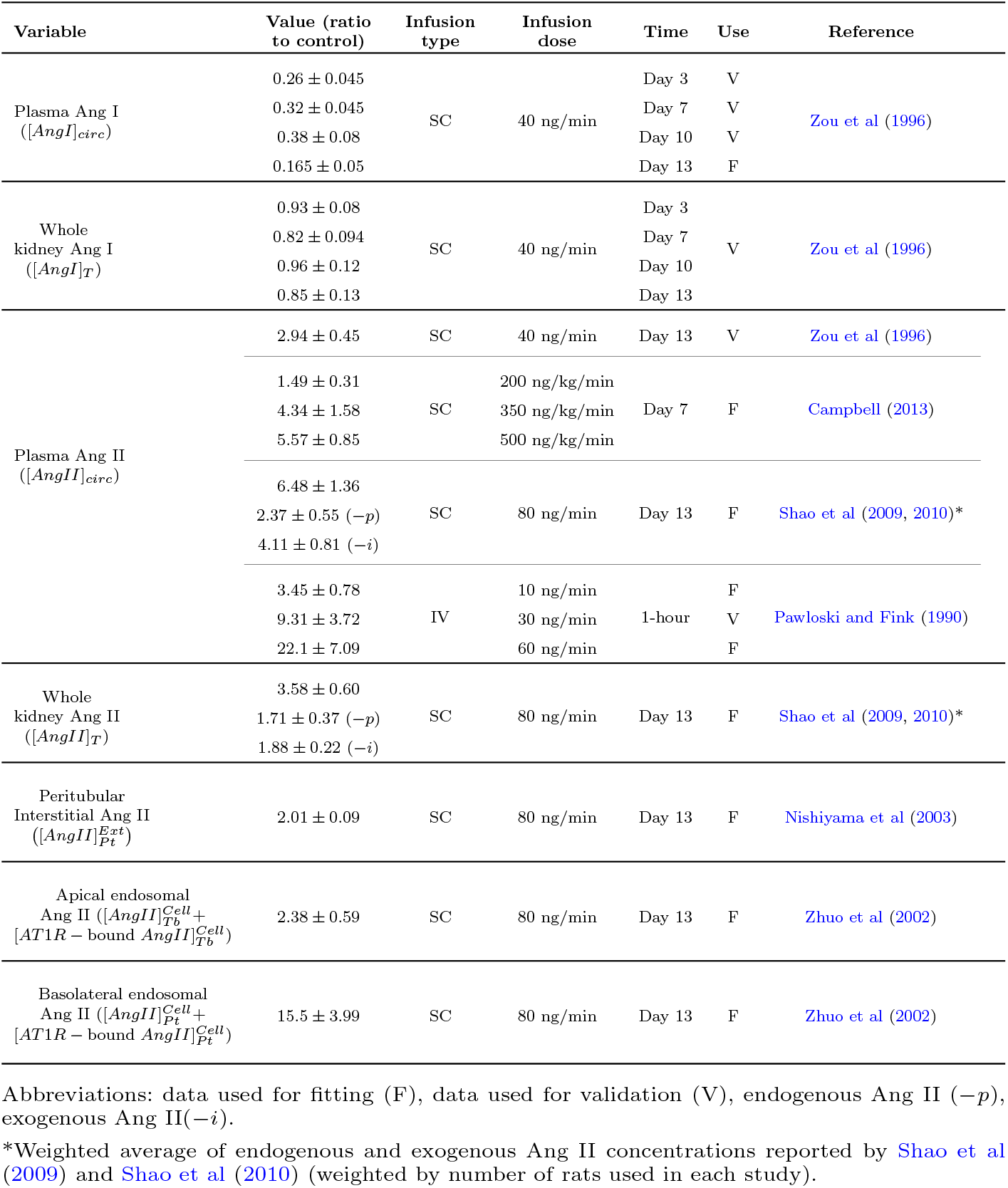
Ang II infusion data used for feedback parameter estimation.

